# ANKS1B in the Nucleus Accumbens controls escalated cocaine self-administration via regulating CBP-FoxO3 complex

**DOI:** 10.64898/2026.01.11.698845

**Authors:** Liping Yang, Xiaoxuan Wu, Xuan Chen, Chao Peng, Zihang Li, Shumin Gao, Shiqiu Meng, Jing Dong, Dong Wu, Liying Lv, Ying Han, Yanxue Xue, Lin Lu, Jie Shi, Jianfeng Liu, Yan Sun

## Abstract

The transition from controlled to escalated drug intake is a core feature of cocaine use disorder (CUD), yet the molecular mechanisms underlying this behavioral escalation remain poorly defined. Our prior genome-wide association study (GWAS) identified *ANKS1B* as a significant shared genetic risk factor for heroin, methamphetamine, and alcohol dependence, suggesting a broad role in addiction vulnerability. However, the specific function of ANKS1B in cocaine addiction and its associated neural mechanisms were unknown. Here, we found that ANKS1B expression level in the Nucleus Accumbens (NAc) was downregulated after extended cocaine use, and manipulating ANKS1B could selectively influence the escalation of cocaine intake and the subsequent cocaine-seeking behavior in the long-access cocaine self-administration rat model. Molecular experiments reveal that ANKS1B interacts with the histone acetyltransferase CBP to control H3K27 acetylation and extended cocaine intake, via epigenetically repressing the transcription factor FoxO3. Overall, these findings suggest that ANKS1B is a crucial factor influencing the escalation of cocaine use. The ANKS1B-CBP-FoxO3 signaling pathway presents a promising target for potential therapeutic interventions for controlling extended cocaine use.

## 1. Introduction

Cocaine use disorder (CUD) continues to be a major public health burden, and no FDA-approved medication is currently available to treat it ^[1, 2]^. A defining feature of CUD is the transition from recreational to compulsive drug use following prolonged cocaine exposure, accompanied by an increase in consumption—a phenomenon termed escalation ^[3]^. Previous research demonstrated that this escalated intake and subsequent compulsive drug-seeking behavior can be reliably modeled using long-access (LgA), but not short-access (ShA), cocaine self-administration paradigms ^[4, 5]^. While research using the LgA model implicates dopaminergic systems and other brain regions in mediating escalated cocaine intake and compulsive use ^[3, 6]^, the molecular mechanisms that underlie escalated cocaine intake remain poorly understood.

Ankyrin repeat domain-containing protein 1B (ANKS1B) is an ankyrin repeat-containing scaffolding protein that regulates synaptic plasticity, glutamatergic signaling, and epigenetically mediated transcriptional programs ^[7–9]^ and has been implicated in various neuropsychiatric disorders ^[7, 10–14]^. ANKS1B exhibits predominant expression in the nucleus accumbens (NAc), where cocaine exposure promotes significant recruitment of chromatin remodeling complexes to its genomic regions ^[15]^. Our previous genome-wide association study (GWAS) identified a common *anks1b* variant (rs2133896) as a shared genetic risk factor for heroin, methamphetamine, and alcohol dependence ^[16, 17]^. In addition, prolonged drug exposure markedly reduces ANKS1B expression, and viral-mediated ANKS1B overexpression in the ventral tegmental area significantly attenuates drug self-administration ^[17]^, suggesting ANKS1B functions as a protective factor against addiction vulnerability.

Extensive research demonstrates that the pathophysiology of CUD is driven by epigenetic regulation, wherein cocaine-induced histone modifications reshape chromatin architecture to induce aberrant gene expression and cellular maladaptations that perpetuate the enduring effects of cocaine exposure ^[18–22]^. Long-term cocaine self-administration leads to persistent increases in histone acetylation within the NAc, a key region mediating reward-related learning ^[23–26]^. As Griffin et al reported, degradation of HDAC4 and HDAC5 in the NAc results in enhanced histone acetylation, thereby facilitating cocaine-induced gene expression and compulsive intake ^[18]^. Previous findings suggested that ANKS1B could be a critical molecular node linking synaptic signaling to epigenetic regulation. Through its ankyrin repeat domains, ANKS1B mediates protein–protein interactions that can couple synaptic activity to chromatin remodeling ^[27, 28]^. Proteins containing ankyrin repeat domains have been shown to interact with histone deacetylases (HDACs) and regulate histone acetylation level ^[29]^, and pathway analyses indicate that ANKS1B deficiency predicts potent inhibition of the NAD⁺-dependent sirtuin signaling cascade (class III HDACs) as well as disruptions in CREB-dependent transcription ^[9]^. Based on this evidence, it might be speculated that ANKS1B could regulate cocaine use via its modulating role in epigenetics in cocaine addiction-related brain regions, such as the NAc.

Here, we report that ANKS1B in the NAc selectively controls escalated cocaine use in the rat LgA cocaine self-administration model. Our findings demonstrate that bidirectional modulation of ANKS1B expression alters the escalation of cocaine consumption and the subsequent incubation of cocaine-seeking behavior. Furthermore, we identify a novel functional axis in which ANKS1B interacts with the histone acetyltransferase CREB-binding protein (CBP) to restrict H3K27 acetylation and suppress transcription of the downstream effector FoxO3 in the NAc. Collectively, these results establish the ANKS1B-CBP-FoxO3 signaling pathway as a critical regulator of escalated cocaine use.

## 2. Results

### 2.1. Long- but not short-access cocaine self-administration reduced ANKS1B expression in the NAc

The *ANKS1B* gene is highly enriched in the NAc compared to other brain regions, as reported by the GTEx consortium (https://gtexportal.org/home/), and its expression is predominantly localized to neurons within the NAc. (Figure S1). To identify the cellular populations expressing ANKS1B protein in the NAc, we conducted immunofluorescence analysis of rat NAc tissue. We found that ANKS1B protein exhibited predominant neuronal localization, as evidenced by robust co-localization with the neuronal marker NeuN and minimal overlap with the astrocytic marker S100β or the microglial marker Iba-1 (Figure S2).

To investigate whether ANKS1B expression correlates with cocaine use patterns, we utilized rodent models of short-access (ShA; 2 h/day) and long-access (LgA; 6 h/day) intravenous cocaine self-administration (**Figure** 1A), which generate distinct trajectories of drug intake^[30, 31]^. As shown in Figure 1B, all animals underwent a 7-day acquisition phase of cocaine self-administration (2 h/day). Following this, ShA rats continued with 10 additional sessions of short-access (2 h/day), whereas LgA rats were switched to 10 sessions of long-access (6 h/day). Notably, only the LgA rats exhibited a progressive escalation in cocaine intake, which is a well-established hallmark of addiction-like behavior ^[4, 32]^. To quantify escalation, we calculated the slope of drug intake escalation across the final 8–17 sessions, which confirmed that only LgA self-administration produced a significant escalation (Figure 1C). Rats were euthanized 24 h after the final self-administration session, and NAc tissue was harvested for Western blot analysis. Results revealed a significant reduction in ANKS1B protein levels in LgA-exposed rats compared to both saline-treated controls and ShA rats (Figure 1D), implicating that ANKS1B expression is negatively associated with escalated cocaine use.

**Figure 1.**
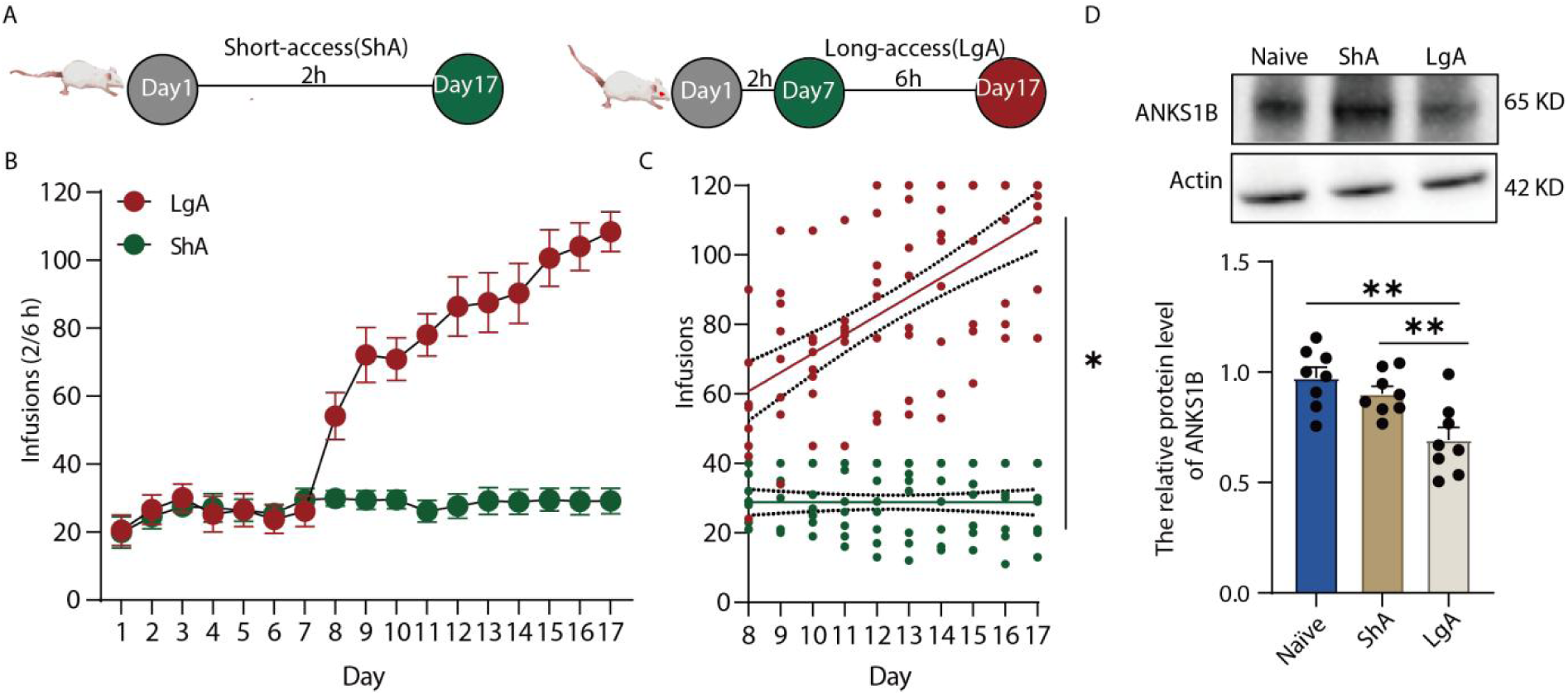
Reduced NAc ANKS1B expression in extended-access rats. (A) Experimental timeline and schematic of the intravenous cocaine self-administration paradigms. Rats were assigned to a short-access (ShA; 2 h/day for 17 days) or a long-access (LgA; 2 h/day for days 1–7, then 6 h/day for days 8–17) schedule. (B) Number of cocaine infusions per session during training for ShA (n = 8) and LgA (n = 8) rats. Data are presented as mean ± SEM. (C) Linear regression analysis of cocaine intake escalation across self-administration sessions in ShA and LgA rats. Each point represents the number of infusions obtained by an individual subject during each session. Solid lines indicate linear regression best-fit lines, and dotted lines indicate 95% confidence intervals. The LgA group showed a significant escalation of cocaine intake across sessions (best-fit line: Y = 5.442X + 17.19, R² = 0.37, F_1,78_ = 46.18, p < 0.01), whereas the ShA group showed no significant escalation (best-fit line: Y = 0.002273X + 28.78, R² = 5.37 × 10⁻⁷, F_1,78_ = 4.19 × 10⁻⁵, p > 0.99). Comparison of regression slopes using an interaction-term model revealed a significant difference between groups (F_1,156_ = 38.70, p < 0.01). (D) Western blot analysis of ANKS1B protein expression in the NAc. Representative blot and quantification for Control (Con), ShA, and LgA groups (n = 8 per group). ANKS1B levels were normalized to actin. One-way ANOVA revealed a significant group effect (F₂, ₂₁ = 9.74, P < 0.01), with post hoc Tukey tests showing increased ANKS1B reduction in LgA compared with Control (P < 0.01) and ShA (P = 0.01). Data are presented as mean ± SEM, P < 0.05 was considered statistically significant. Statistical significance was determined by one-way ANOVA followed by Tukey’s post hoc test.

### 2.2. Viral-mediated manipulation of ANKS1B expression in the NAc bidirectionally regulated escalated cocaine intake and the subsequent cocaine-seeking behavior

To investigate whether ANKS1B in the NAc plays a critical role in the development of extended cocaine use, we employed adeno-associated virus (AAV)-mediated gene delivery to modulate ANKS1B expression within the NAc and assessed its impact on cocaine self-administration behavior (**Figure** 2A). Immunofluorescence and Western blot analyses confirmed the successful overexpression or knockdown of ANKS1B following AAV-*Anks1b*-Flag or AAV-*Anks1b*-shRNA-mCherry delivery, respectively (Figure 2B, C, K, L). Behavioral experiments revealed that *Anks1b* overexpression in the NAc did not alter cocaine self-administration acquisition during ShA sessions (Figure 2D, E). Notably, whereas control rats exhibited a progressive increase in cocaine intake over LgA sessions, ANKS1B overexpressing (OE) rats maintained a stable number of infusions throughout all LgA sessions (Figure 2F). A significant difference between control and OE rats was observed on the final (day 16–17), but not the initial (day 8–9), LgA session (Figure 2G). Consistently, ANKS1B overexpression also reduced cue-induced cocaine-seeking behavior following 1 or 28 days of abstinence from cocaine (Figure 2H, I). To address the concern regarding neuronal specificity, we used a neuron-specific lentiviral vector (LV-hsyn-Anks1b) to overexpress ANKS1B in the NAc. This manipulation recapitulated the AAV-mediated effects by attenuating cocaine intake escalation and cue-induced seeking (Figure S3).

**Figure 2:**
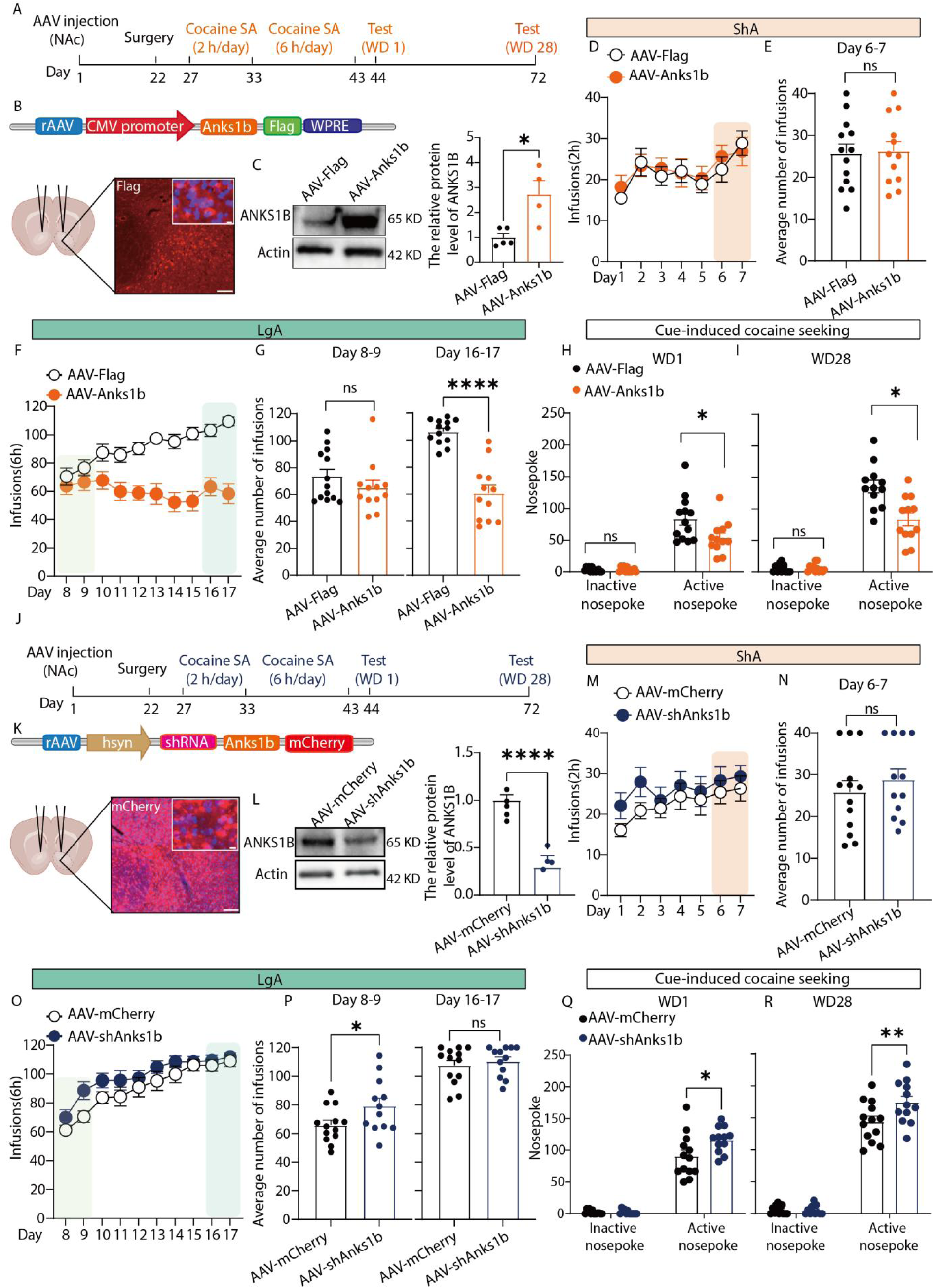
ANKS1B in the nucleus accumbens (NAc) bidirectionally regulates cocaine self-administration and seeking. (A) Experimental timeline showing AAV injection, cocaine self-administration, and withdrawal (WD) testing. (B) Representative image showing Flag-tagged ANKS1B expression (red) in the NAc. Scale bars, 100 μm and 10 μm. (C) Western blot validation confirming successful ANKS1B overexpression (n = 4) in the NAc compared with the AAV-Flag group (n = 5) (unpaired t-test, t₇ = 3.31, P = 0.01). (D) ANKS1B overexpression did not alter cocaine acquisition during the ShA phase (n = 13 and 12 for the AAV-Flag and AAV-Anks1b groups). (E) Comparison of cocaine infusions during the last 2 days of ShA (days 6–7) revealed no difference between groups (unpaired t-test, t₂_3_ = 0.17, P = 0.87). (F) ANKS1B overexpression prevented escalation of cocaine intake during the LgA phase. (G) Comparison of infusions on the first 2 days (days 8–9; unpaired t-test, t₂_3_ = 1.12, P = 0.27) and the last 2 days (days 16–17; t₂_3_ = 7.18, P < 0.01) of LgA. (H and I) Overexpression of ANKS1B significantly reduced cue-induced cocaine seeking (Active nosepoke) on both WD1 and WD28. (WD1: two-way RM ANOVA, genotype × nosepoke interactions: F_1, 23_ = 5.14, P = 0.03; post hoc, P < 0.01; WD28: two-way RM ANOVA, genotype × nosepoke interactions: F_1, 22_ = 12.56, P < 0.01; post hoc, P < 0.01; n = 12–13 per group). (J) Experimental design for shRNA-mediated ANKS1B knockdown (KD). (K and L) Validation of ANKS1B KD by mCherry fluorescence (K) and Western blot (L) (unpaired t-test, t₇ = 8.50, P < 0.01; n = 4–5 per group). (M) ANKS1B knockdown did not affect cocaine acquisition during the ShA phase (n = 13 and 12 for the AAV-mCherry and AAV-shAnks1b-mCherry). (N) Comparison of cocaine infusions during the last 2 days of ShA (days 6–7) revealed no group difference (unpaired t-test, t_23_ = 0.78, P = 0.44). (O) ANKS1B knockdown potentiated the escalation of cocaine intake during the LgA phase. (P) Comparison of infusions on the first 2 days (days 8–9; unpaired t-test, t_23_ = 2.11, P = 0.04) and the last 2 days (days 16–17; unpaired t-test, t₂_3_ = 0.66, P = 0.52) of LgA. (Q and R) Knockdown of ANKS1B significantly increased cue-induced cocaine seeking (Active nosepoke) on both WD1 and WD28. (WD1: two-way RM ANOVA, genotype × nosepoke interactions: F_1, 23_ = 5.24, P = 0.03; post hoc, P < 0.01. WD28: two-way RM ANOVA, genotype × nosepoke interactions: F_1, 23_ = 5.95, P = 0.02; post hoc, P < 0.01, n = 13 and 12 for the AAV-mCherry and AAV-shAnks1b-mCherry). Data are presented as mean ± SEM. P < 0.05 was considered statistically significant; ns, not significant. Average number of infusions: Each symbol represents one animal’s average infusions over two consecutive sessions. Statistical significance was determined by two-way repeated-measures (RM) ANOVA followed by Bonferroni’s multiple comparisons test.

In contrast, virus-mediated knockdown of ANKS1B accelerated the escalation of cocaine intake during LgA sessions without affecting acquisition during ShA sessions (Figure 2J–P). ANKS1B knockdown also enhanced cue-induced cocaine-seeking behavior after abstinence (Figure 2Q, R). To determine whether ANKS1B manipulation affects the maintenance of ShA cocaine intake, we employed the same viral gene expression approach and evaluated its impact over a 17-day ShA self-administration period. Neither ANKS1B knockdown nor overexpression altered stable cocaine intake when animals were maintained on a ShA schedule for 17 consecutive days (Figure S4).

Most neurons in the NAc express dopamine receptors D1 or D2, comprising two primary cell types: D1- and D2-expressing medium spiny neurons (MSNs). Single-cell RNA sequencing data ^[33]^ and anatomical analyses revealed that ANKS1B is expressed in both D1- and D2-MSNs (Figure S5A-D). To assess whether ANKS1B in these distinct cell populations contributes differentially to cocaine escalation, we injected AAV-D1-Cre-EGFP or AAV-D2-Cre-mCherry into *Anks1b* double-floxed (Anks1b^f/f^) rats to achieve cell-type-specific deletion of *Anks1b* in D1- or D2-MSNs. However, both groups exhibited increased cue-induced cocaine-seeking behavior, indicating that ANKS1B in both neuronal subtypes is required for regulating relapse-like behavior rather than cocaine intake escalation (Figure S6). Collectively, these results demonstrate that NAc ANKS1B negatively regulates the escalation of cocaine use.

### 2.3. ANKS1B modulation in the NAc did not lead to nonspecific behavioral suppression or influence sucrose self-administration

To rule out nonspecific effects of ANKS1B manipulation on locomotion, anxiety, or natural reward processing, we evaluated the consequences of NAc ANKS1B overexpression or knockdown in the open field test (OFT), elevated plus maze (EPM), novel object recognition (NOR), and sucrose self-administration. ANKS1B overexpression or knockdown did not alter locomotor activity or center exploration time in the OFT (**Figure** 3A–D), time spent in open or closed arms in the EPM (Figure 3E–H), or recognition memory performance in the NOR task (Figure 3I–L). Additionally, ANKS1B manipulation did not affect ShA sucrose self-administration (days 1–7), LgA sessions (days 8–17), or cue-induced sucrose-seeking after 1 or 28 days of abstinence (Figure 3M–V). Together, these findings confirm that NAc ANKS1B regulation selectively influences cocaine-related behaviors without inducing general behavioral suppression or altering natural reward intake or motivation.

**Figure 3.**
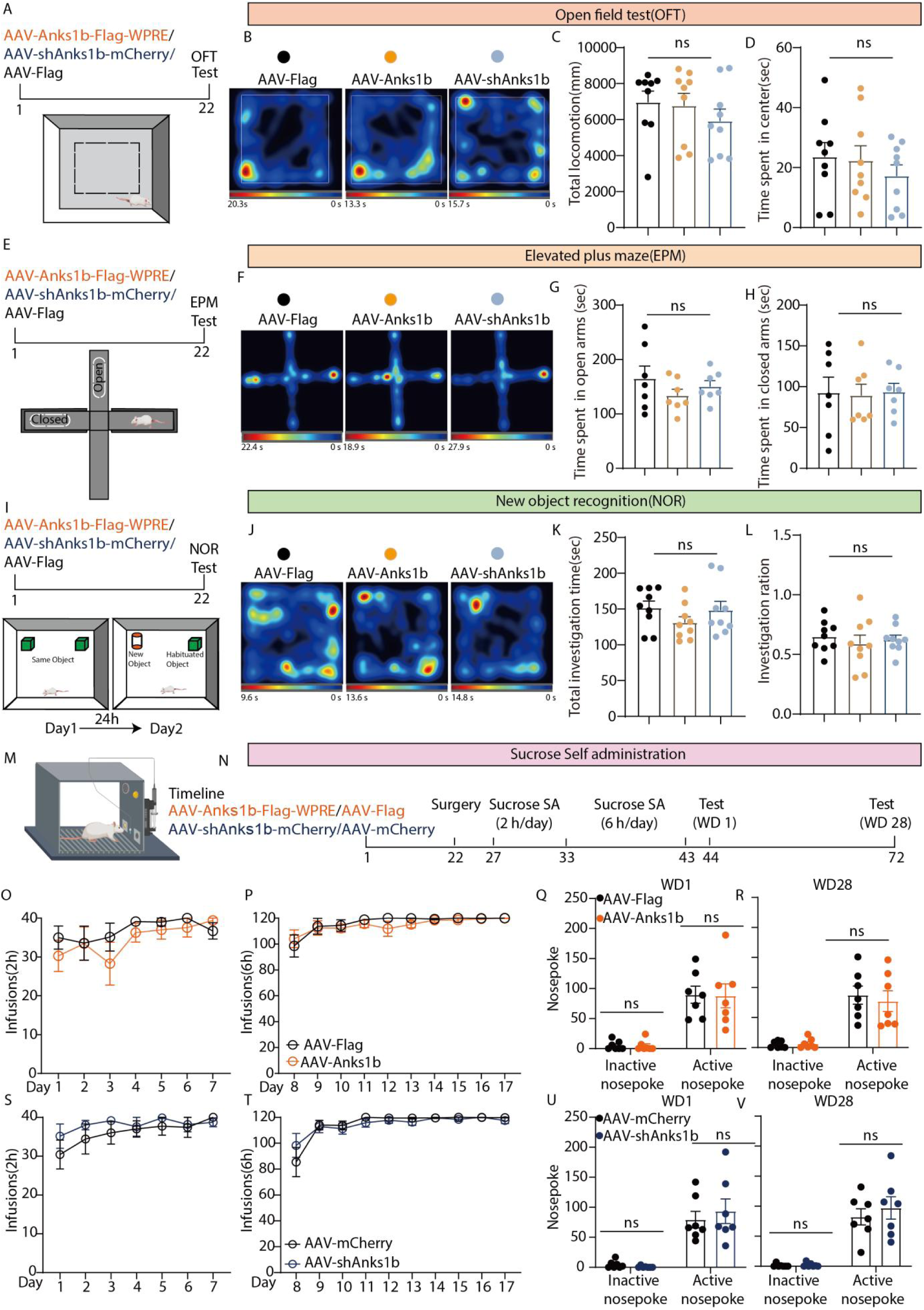
ANKS1B manipulation in the NAc does not affect general locomotion, anxiety, recognition memory, or natural reward seeking. (A–D) Open field test (OFT). Neither overexpression (OE) nor knockdown (KD) of ANKS1B in the NAc altered total locomotion (one-way ANOVA, F_2, 24_ = 0.72, P = 0.50) or time spent in the center (one-way ANOVA, F₂,₂₄ = 0.56, P = 0.58; n = 9 per group). (E–H) Elevated plus maze (EPM). ANKS1B OE and KD did not affect time spent in the open arms (one-way ANOVA, F₂,₁₈ = 0.02, P = 0.98) or closed arms (one-way ANOVA, F₂,₁₈ = 0.98, P = 0.40; n = 7 per group). (I–L) Novel object recognition (NOR) test. ANKS1B OE and KD did not affect total object investigation time (one-way ANOVA, F₂,₂₄ = 1.22, P = 0.31) or the novel object discrimination ratio (one-way ANOVA, F₂,₂₄ = 0.33, P = 0.72; n = 9 per group). (M–V) Sucrose self-administration. ANKS1B OE and KD did not alter the acquisition of sucrose self-administration (O, P, S, T) or cue-induced sucrose seeking (Active nosepoke) (Q, R, U, V). (OE WD1: two-way RM ANOVA, genotype × nosepoke interactions: F_1, 12_ = 0.01, P = 0.96; OE WD28: two-way RM ANOVA, genotype × nosepoke interactions: F_1, 12_ = 0.29, P = 0.60; KD WD1: two-way RM ANOVA, genotype × nosepoke interactions: F_1, 12_ = 0.53, P = 0.48; KD WD28: two-way RM ANOVA, genotype × nosepoke interactions: F_1, 12_ = 0.40, P = 0.54; n = 7 per group). Heat maps illustrate the spatial distribution of exploratory activity in the open field test. Heat maps illustrate the spatial distribution of exploratory activity in the open field test. Warmer colors indicate longer residence time, whereas cooler colors indicate lower occupancy. The color scale represents the relative time spent in each area of the arena. Data are presented as mean ± SEM. P < 0.05 was considered statistically significant.

### 2.4. ANKS1B overexpression attenuated long-access cocaine intake-induced structural and synaptic plasticity

Cocaine addiction has been closely linked to synaptic plasticity ^[34–36]^. To investigate the structural changes in NAc neurons following cocaine exposure, we utilized Golgi staining. This approach also allowed us to evaluate the effects of ANKS1B overexpression on cocaine-induced morphological alterations. Compared to rats under ShA cocaine self-administration, those under LgA conditions exhibited a significant increase in total dendritic spine density. Importantly, overexpression effectively prevented this LgA cocaine-induced elevation in spine density (**Figure 4**A, B). Further subtype analysis revealed that LgA cocaine preferentially increased the density of thin and mushroom spines, with no significant effects on stubby or filopodia spines. Notably, ANKS1B overexpression selectively reversed the increase in thin spines induced by LgA cocaine, while showing only a modest influence on the mushroom spine population (Figure 4C–F).

**Figure 4.**
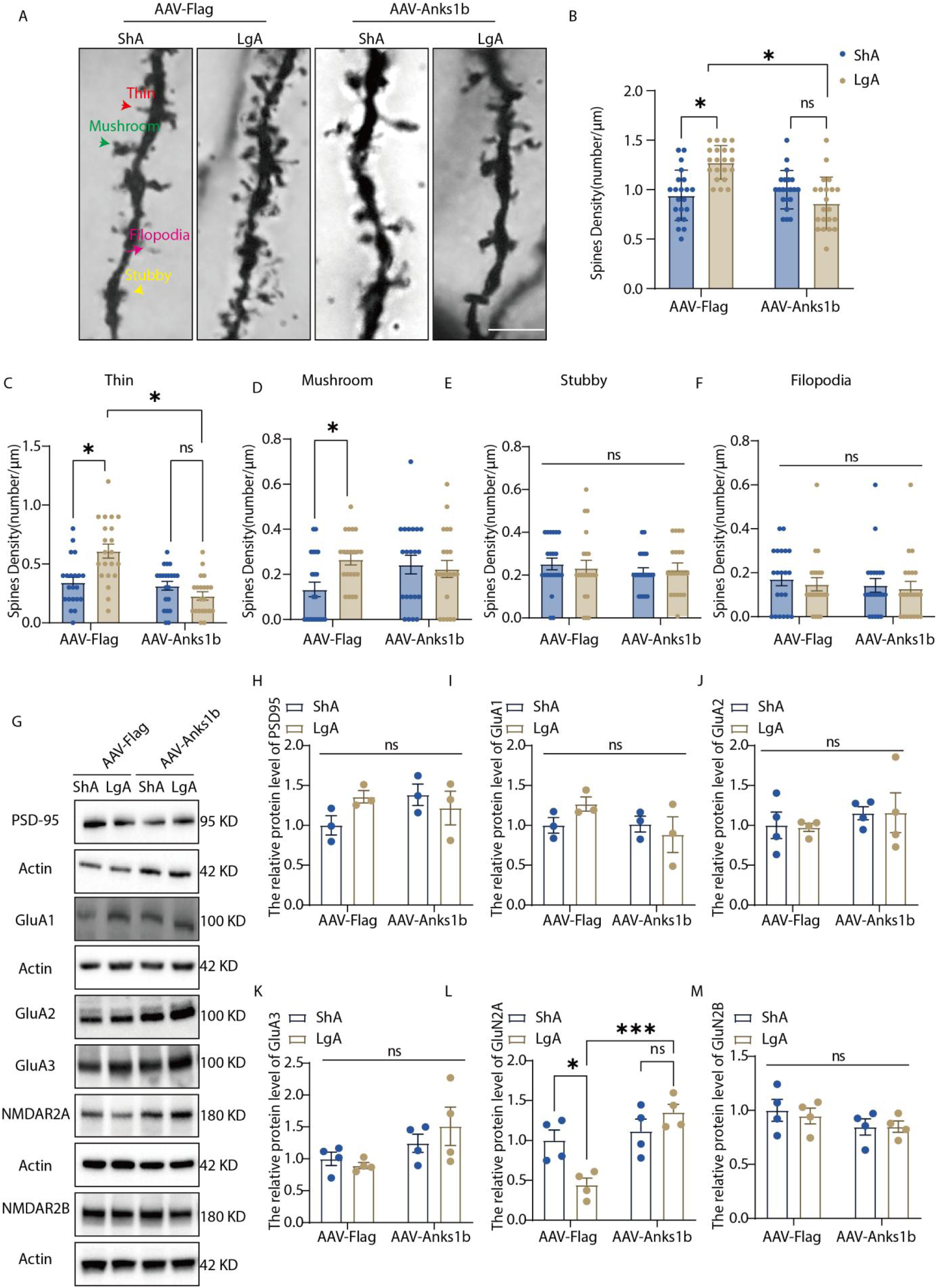
ANKS1B overexpression in the NAc prevents cocaine-induced structural and molecular synaptic plasticity. (A) Representative Golgi-stained images of medium spiny neuron dendrites in the NAc from AAV-Flag and AAV-Anks1b rats following either ShA or LgA cocaine self-administration. Arrows indicate representative thin (red), mushroom(green), stubby (yellow), and filopodia (pink) spine morphologies. (B–F) LgA cocaine self-administration induced a significant increase in the density of total dendritic spines (B), which was driven by an increase in both thin (C) and mushroom (D) spines. ANKS1B OE completely blocked this effect. Stubby (E) and filopodia (F) spine densities were unaffected by either cocaine history or genotype (two-way ANOVA, Total: genotype × access interaction: F_1, 80_ = 23.11, P < 0.01; post hoc, AAV-Flag-ShA vs. AAV-Flag-LgA, P < 0.01; AAV-Flag-LgA vs. AAV-Anks1b-LgA, P < 0.01; Thin: genotype × access interaction: F_1, 80_ = 15.17, P = 0.0002; post hoc, AAV-Flag-ShA vs. AAV-Flag-LgA, P = 0.0004; AAV-Flag-LgA vs. AAV-Anks1b-LgA, P < 0.01; Mushroom: genotype × access interaction: F_1, 80_ = 4.82, P = 0.03 post hoc, AAV-Flag-ShA vs. AAV-Flag-LgA, P = 0.03; AAV-Flag-LgA vs. AAV-Anks1b-LgA, P = 0.82;. Stubby: genotype × access interaction: F_1, 80_ = 0.27, P = 0.61. Filopodia: genotype × access interaction: F_1, 80_ = 0.02, P = 0.89; n = 5 rats per group). (G) Representative Western blots of synaptic proteins from NAc tissue. (H–M) Quantification of protein levels revealed no significant changes in PSD-95, GluA1, GluA2, GluA3, or NR2B across groups(two-way ANOVA, PSD-95: F_1, 8_ = 3.33, P = 0.11; GluA1: F_1, 8_ = 2.04, P = 0.19; GluA2: F_1, 12_ = 0.01; P = 0.92; GluA3: F_1, 12_ = 1.12, P = 0.31; NR2B: F_1, 12_ = 0.13; P = 0.73). In contrast, LgA significantly decreased NR2A protein levels (L), an effect that was reversed by ANKS1B OE (NR2A: F_1, 12_ = 10.97, P < 0.01; post hoc, AAV-Flag-LgA vs. AAV-Flag-ShA, P = 0.03; AAV-Anks1b-LgA vs. AAV-Flag-LgA, P < 0.01; n = 3–4 per group). Data are presented as mean ± SEM. P < 0.05 was considered statistically significant. Statistical significance was determined by two-way ANOVA followed by Tukey’s post hoc test.

To explore whether these structural changes were accompanied by molecular alterations associated with synaptic plasticity, we isolated postsynaptic membrane fractions from NAc tissue following ShA or LgA cocaine self-administration and performed Western blot analyses. Among the examined synaptic proteins—including AMPA receptor subunits (GluA1–3), NMDA receptor subunits (NR2A and NR2B), and PSD-95—we observed a significant reduction in membrane-localized NR2A expression following LgA cocaine exposure. This decrease was prevented by ANKS1B overexpression (Figure 4K–M). Taken together, these findings indicate that modulation of ANKS1B can prevent the structural and molecular remodeling of synaptic components in the NAc that are associated with extended cocaine intake.

### 2.5. ANKS1B interacts with the histone acetyltransferase CBP to control H3K27 acetylation and extended cocaine intake

Previous studies have suggested an interaction between ANKS1B and CBP, a transcriptional co-activator endowed with histone acetyltransferase (HAT) activity ^[37–40]^. Given the established role of histone acetylation-mediated epigenetic regulation in the NAc in modulating compulsive cocaine self-administration ^[18, 41]^, we hypothesized that ANKS1B may influence the development of extended cocaine intake through the modulation of histone acetylation dynamics mediated by its interaction with CBP. To test this possibility, we performed co-immunoprecipitation (Co-IP) experiments using NAc tissue lysates, which demonstrated a physical interaction between endogenous ANKS1B and CBP (**Figure 5**A). This interaction was further supported by immunofluorescence imaging, which revealed overlapping nuclear localization patterns of ANKS1B and CBP (Figure 5B). Importantly, ANKS1B-CBP binding was markedly reduced in LgA compared with ShA rats, whereas ANKS1B overexpression restored this interaction (Figure 5C, D). Importantly, CBP protein levels were not affected by cocaine exposure or ANKS1B overexpression (Figure S7), suggesting that the reduced interaction is independent of CBP expression.

**Figure 5.**
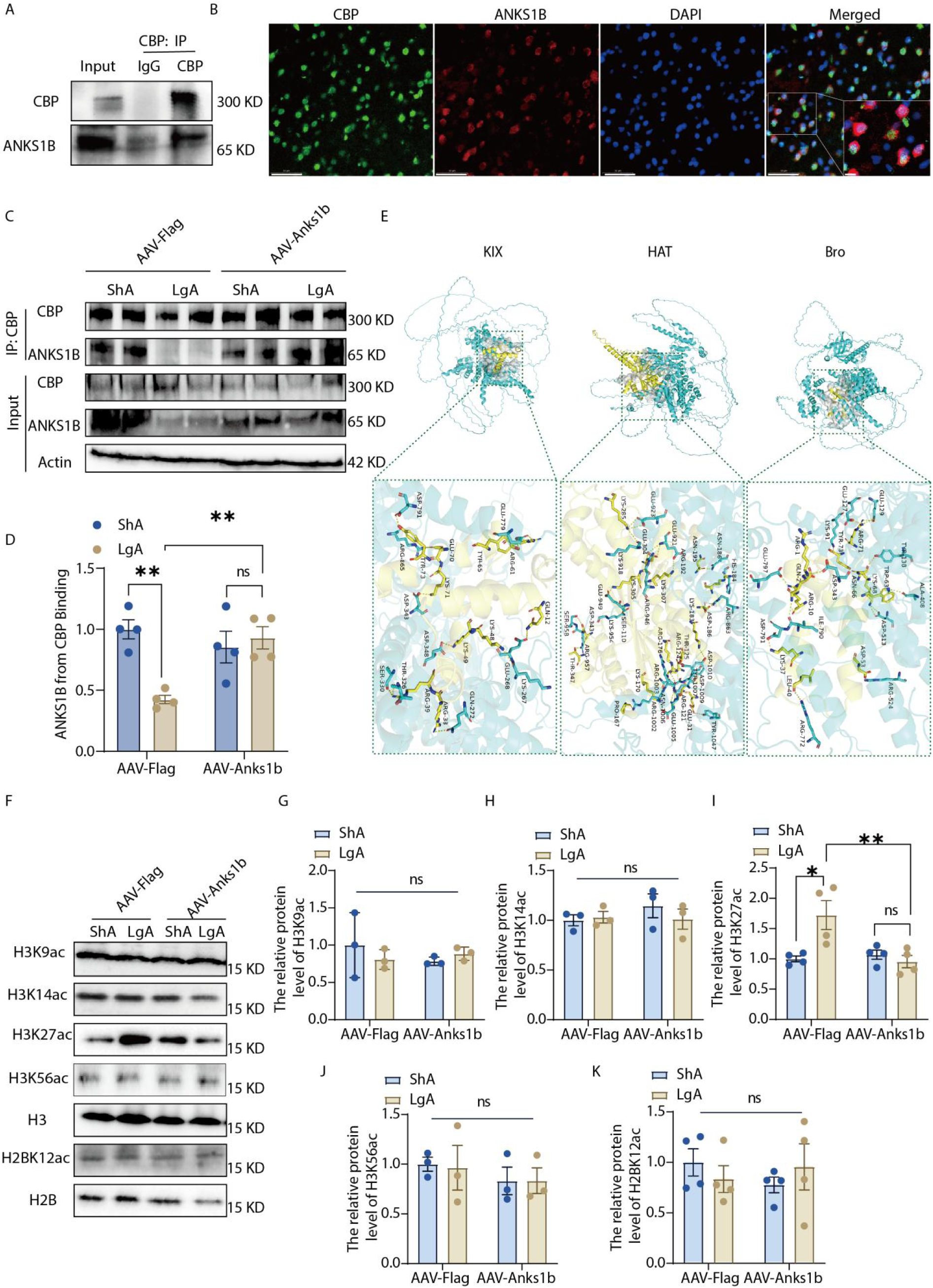
ANKS1B interacts with the histone acetyltransferase CBP to regulate H3K27 acetylation. (A) Co-immunoprecipitation (Co-IP) of NAc lysates using an anti-CBP antibody successfully pulled down ANKS1B, confirming their interaction. (B) Confocal images showing co-localization (yellow, arrows) of CBP (green) and ANKS1B (red). Scale bar, 50 µm and 5 µm. (C) Representative Co-IP and input western blots showing ANKS1B-CBP binding in ShA and LgA rats with or without ANKS1B overexpression. (D) Quantification of ANKS1B-CBP binding. LgA significantly reduced ANKS1B-CBP interaction compared with ShA, whereas ANKS1B overexpression restored this interaction (two-way ANOVA, genotype × access interaction: F_1, 12_ = 13.10, P < 0.01; post hoc, AAV-Flag-ShA vs. AAV-Flag-LgA, P < 0.01, AAV-Flag-LgA vs. AAV-Anks1b-LgA, P < 0.01). (E) Molecular docking simulation showing binding interfaces between ANKS1B (yellow) and the Bromodomain (Bro), Histone Acetyltransferase (HAT), and KIX domains of CBP (cyan). (F) Representative Western blots for total and acetylated histone H3 and H2B. (G–K) Quantification of histone acetylation levels. LgA increased H3K27ac levels, which were restored by ANKS1B OE (two-way ANOVA, genotype × access interaction: F_1, 12_ = 9.29, P = 0.01; post hoc, AAV-Flag-ShA vs. AAV-Flag-LgA, P = 0.01; AAV-Flag-LgA vs. AAV-Anks1b-LgA, P = 0.01). No changes were observed for H3K9ac (genotype × access interaction: F_1, 8_ = 1.16, P = 0.31), H3K14ac (genotype × access interaction: F_1, 8_ = 0.87, P = 0.38), H3K56ac (genotype × access interaction: F_1, 8_ = 0.01, P = 0.90), or H2BK12ac (genotype × access interaction: F_1, 12_ = 1.26, P = 0.28); n = 3–4 per group. Data are presented as mean ± SEM. P < 0.05 was considered statistically significant. Statistical significance was determined by two-way ANOVA followed by Tukey’s post hoc test.

CBP contains several major domains, including KIX, bromodomain (Bro), and HAT, which are required for robust acetyltransferase activity in vitro and in vivo. These domains control either the interaction with different proteins or the acetyltransferase activity of CBP ^[42, 43]^. Our computational protein–protein docking analysis revealed a predicted stable interface between ANKS1B and the BRO/KIX/HAT domains of CBP (Figure 5E). Predicted aligned error (PAE) and per-residue confidence (pLDDT) analyses indicated reliable structural modeling of ANKS1B interactions with the CBP BRO, HAT, and KIX domains, supporting the stability of the predicted interfaces (Figure S8). Western blot analysis revealed that LgA cocaine exposure selectively increased H3K27ac levels in the NAc compared with ShA rats (Figure 5F, I). Notably, this effect was completely prevented by ANKS1B overexpression (OE-LgA), indicating that ANKS1B restrains CBP-mediated H3K27 acetylation. Other acetylation marks, including H3K9ac, H3K14ac, H3K56ac, and H2BK12ac, remained unchanged (Figure 5G–H, J–K).

To directly test whether CBP activity mediates cocaine-related behaviors, we inhibited CBP in the NAc with C646, a CBP HAT domain-specific inhibitor that can block H3K27ac activity ^[44–46]^. Our results showed that intra-NAc C646 infusion did not alter cocaine acquisition under ShA conditions, but significantly reduced escalation during LgA and suppressed cue-induced cocaine seeking after 1- and 28-day abstinence, an effect resembling those of ANKS1B overexpression (Figure S9). These findings suggest that ANKS1B acts as an inhibitory scaffold for CBP to regulate extended cocaine intake.

### 2.6. ANKS1B controls LgA cocaine intake-induced FoxO3 transcription

To define ANKS1B-dependent transcriptional regulation in the nucleus accumbens, we performed RNA sequencing. Using a consistent comparison framework, long-access cocaine exposure (LgA vs ShA) induced substantial transcriptional changes, including 1,717 upregulated genes, indicating transcriptional activation associated with escalated cocaine intake. Among these, 604 genes that were upregulated under LgA (relative to ShA) were subsequently downregulated upon ANKS1B overexpression. An additional 41 genes exhibited the opposite pattern, being downregulated under LgA and upregulated upon ANKS1B overexpression. Together, these 645 genes constitute the full set of transcripts whose expression is reversed by ANKS1B (**Figure 6**A; Figure S10). Heatmap visualization based on these 645 genes showed that LgA induced increases in gene expression, which were largely restored toward baseline levels by ANKS1B overexpression (Figure 6B). KEGG pathway analysis of the 645 reversed genes identified several significantly enriched pathways, with the FoxO signaling pathway emerging as a dominant regulatory node (Figure 6C). Previous studies have implicated a role of FoxO transcription factors in synaptic plasticity, stress responses, and addiction-related neuroadaptations ^[47–49]^. Then, we focused on this pathway for further analysis.

**Figure 6.**
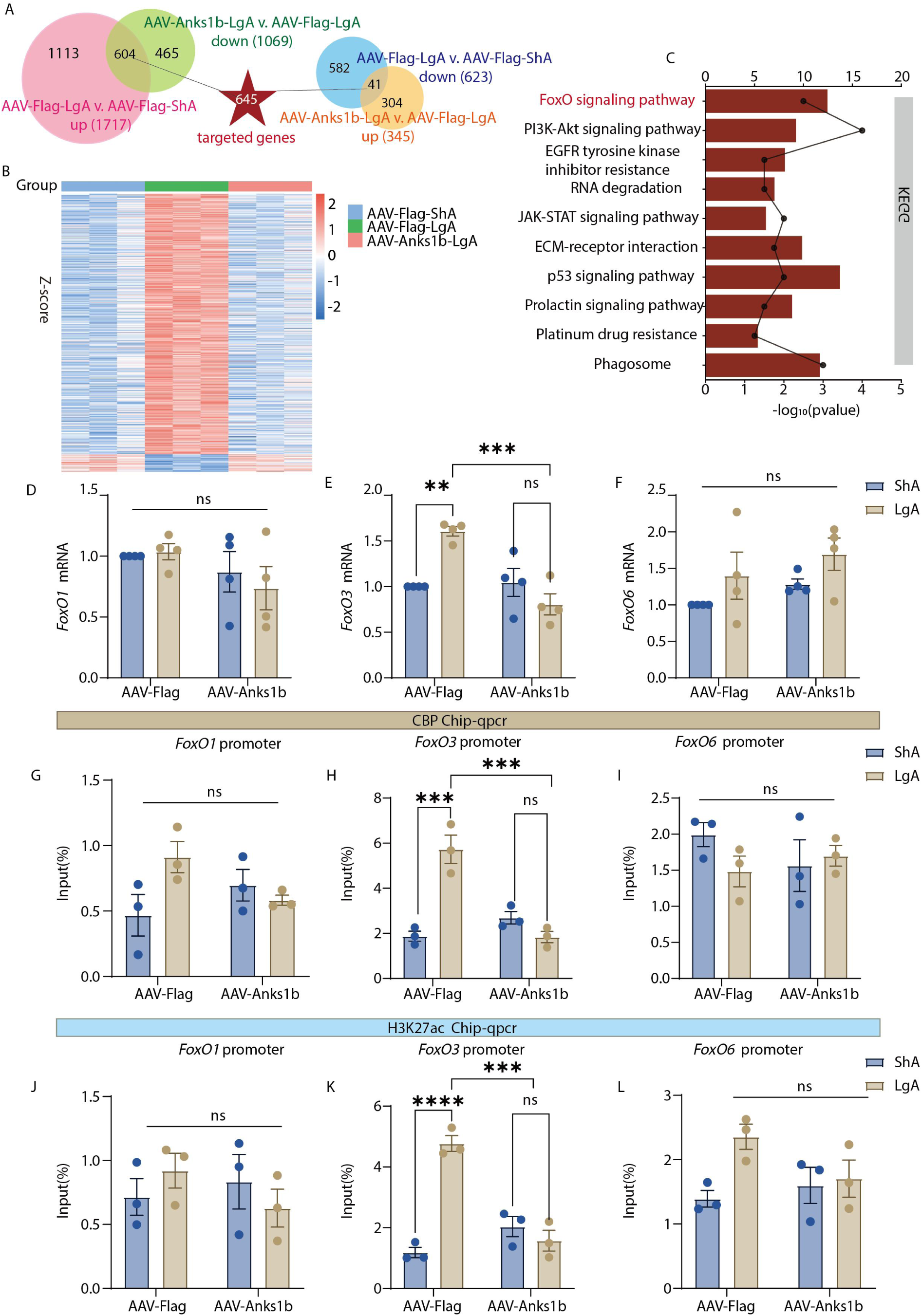
The ANKS1B-CBP complex epigenetically regulates the FoxO signaling pathway by targeting the FoxO3 promoter. (A) Venn diagram of differentially expressed genes. (B) Each row of the heatmap depicts the Z-score-transformed log_2_(1+FPKM) expression values for an individual differentially expressed gene across all samples (blue indicates low expression, red indicates high expression). (C) KEGG pathway analysis of overlapping genes. (D–F) FoxO1 mRNA, FoxO3 mRNA and FoxO6 mRNA (two-way ANOVA, FoxO1: genotype × access interaction: F_1, 12_ = 0.46, P = 0.51; FoxO3: genotype × access interaction: F_1, 12_ = 18.43, P < 0.01; post hoc, AAV-Flag-ShA vs. AAV-Flag-LgA, P = 0.01; AAV-Flag-LgA vs. AAV-Anks1b-LgA, P = 0.01; FoxO6: genotype × access interaction: F_1, 12_ = 0.01, P = 0.98; n = 4 rats per group). (G–I) ChIP-qPCR for CBP at FoxO1, FoxO3 and FoxO6 promoter (two-way ANOVA, FoxO1: genotype × access interaction: F_1, 8_ = 5.66, P = 0.04; post hoc, AAV-Flag-ShA vs. AAV-Flag-LgA, P = 0.1; AAV-Flag-LgA vs. AAV-Anks1b-LgA, P = 0.27; FoxO3: genotype × access interaction: F_1, 8_ = 38.02, P < 0.01; post hoc, AAV-Flag-ShA vs. AAV-Flag-LgA, P < 0.01; AAV-Flag-LgA vs. AAV-Anks1b-LgA, P < 0.01; FoxO6: genotype × access interaction: F_1, 8_ = 1.88, P = 0.21; n = 3 rats per group). (J–L) ChIP-qPCR for H3K27ac at FoxO1, FoxO3 and FoxO6 promoter (two-way ANOVA, FoxO1: genotype × access interaction: F_1, 8_ = 1.60, P = 0.24; FoxO3: genotype × access interaction: F_1, 8_ = 50.65, P < 0.01; post hoc, AAV-Flag-ShA vs. AAV-Flag-LgA, P < 0.01; AAV-Flag-LgA vs. AAV-Anks1b-LgA, P < 0.01. FoxO6: genotype × access interaction: F_1, 8_ = 3.39, P = 0.10; n = 3 rats per group). Data are presented as mean ± SEM. P < 0.05 was considered statistically significant. Statistical significance was determined by two-way ANOVA followed by Tukey’s post hoc test.

Our qPCR analyses revealed a robust upregulation of *FoxO3* mRNA in LgA rats, an effect completely abolished by ANKS1B overexpression (Figure 6E). In contrast, other *FoxO* subtypes (*FoxO1* and *FoxO6*) showed no significant changes in association with ANKS1B overexpression (Figure 6D, F). There is evidence that *FoxO* expression could be regulated by histone acetylation^[50]^; we selected this pathway for focused mechanistic investigation. We next investigated whether *FoxO3* expression is regulated at the chromatin level using chromatin immunoprecipitation followed by quantitative PCR (ChIP-qPCR). We observed a significant increase in CBP occupancy and H3K27ac deposition at the *FoxO3* promoter in LgA rats, indicative of enhanced transcriptional activation. More importantly, ANKS1B overexpression completely abolished both CBP recruitment and H3K27ac enrichment (Figure 6H, K). In contrast, neither CBP binding nor H3K27ac levels at the *FoxO1* or *FoxO6* promoters were affected by ANKS1B overexpression (Figure 6G, I, J, L). Together, these results indicate that ANKS1B controls LgA cocaine intake-evoked *FoxO3* transcription, likely by regulating CBP recruitment and H3K27ac deposition at its promoter.

### 2.7. FoxO3 is a necessary downstream mediator of the ANKS1B pathway for cocaine-related behaviors

To determine whether FoxO3 mediates ANKS1B’s role in regulating escalated cocaine use, we assessed the impact of FoxO3 manipulation on ANKS1B overexpression. We bilaterally injected viruses to co-express ANKS1B and *FoxO3* in the NAc (similar to Figure 2A; **Figure 7**A) before behavioral testing. Fluorescence microscopy and Western blotting confirmed successful *FoxO3* overexpression (Figure 7B, C). Consistent with prior results, ANKS1B overexpression alone prevented cocaine intake escalation (Figure 7E, F). Notably, co-expression of *FoxO3* reversed this protective effect, restoring escalated intake. Similar results were observed during cue-induced seeking after abstinence (Figure 7G, H). These findings identify FoxO3 as a critical downstream effector of ANKS1B in driving cocaine intake escalation. To further validate FoxO3’s role in mediating cocaine intake escalation, we pharmacologically inhibited the FoxO pathway using carbenoxolone, a selective FoxO3 inhibitor with demonstrated high potency (Figure 7I)^[51, 52]^. Intra-NAc injection of carbenoxolone significantly reduced cocaine intake during LgA (but not ShA) (Figure 7J, K) and attenuated drug-seeking behavior during abstinence (Figure 7L, M). Collectively, these results establish FoxO3 as a necessary downstream effector for ANKS1B overexpression’s inhibitory effects on escalated cocaine consumption.

**Figure 7.**
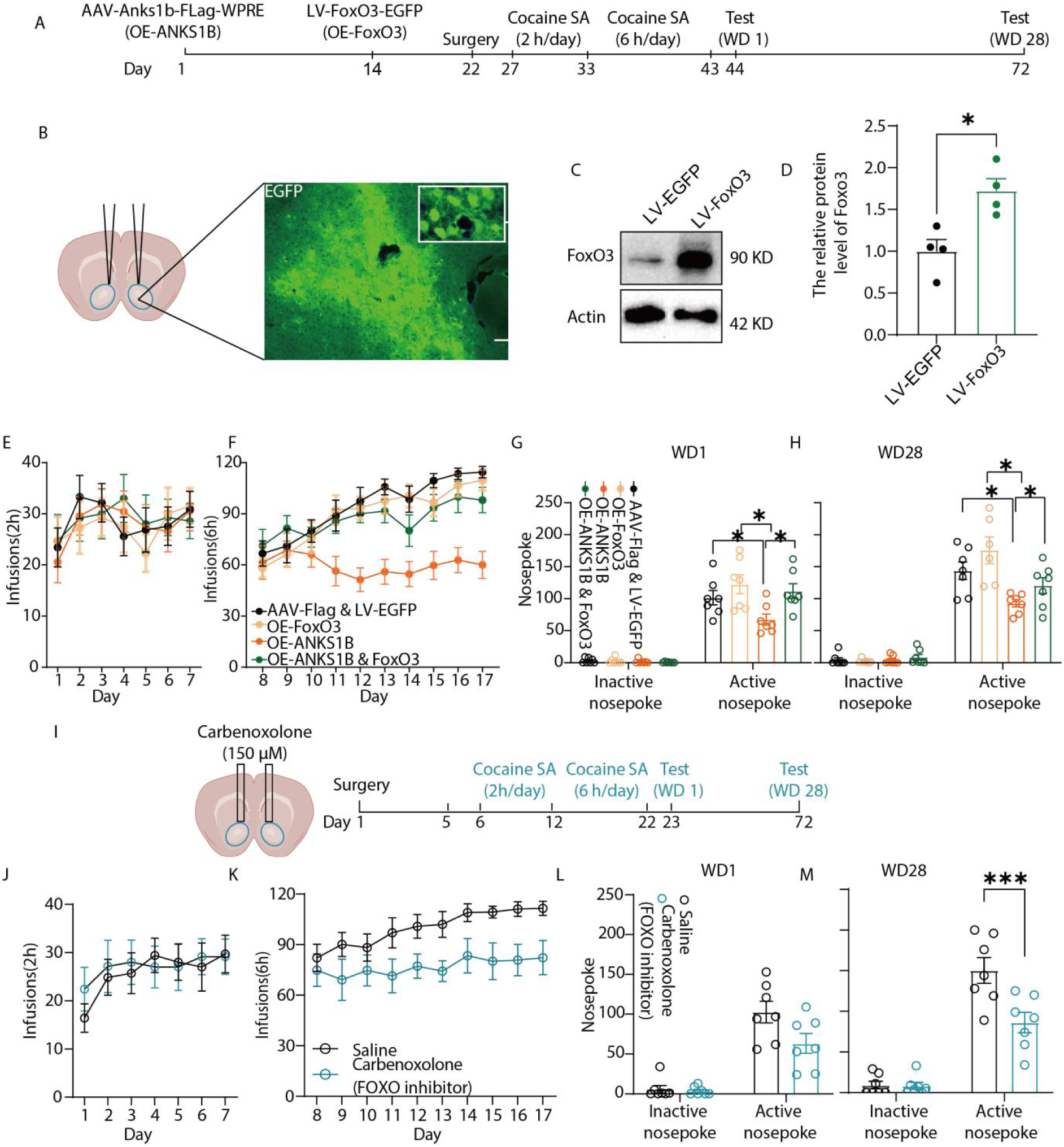
FoxO3 in the NAc is a necessary and sufficient mediator of cocaine-seeking behaviors. (A) Experimental timeline of ANKS1B and FoxO3 overexpression in the NAc followed by behavioral testing. (B) Representative EGFP fluorescence confirming viral transduction. Scale bars, 100 µm and 50 µm. (C–D) Western blot validation showing increased FoxO3 protein levels (unpaired t test, t_6_ = 3.55, P = 0.01; n = 4 per group). (E) FoxO3 overexpression did not affect cocaine acquisition during the ShA phase. (F) During the LgA phase, co-overexpression of FoxO3 with ANKS1B reversed the protective effect of ANKS1B OE, resulting in an escalation of cocaine intake similar to controls. (G–H) Co-overexpression of FoxO3 also abolished the suppressive effect of ANKS1B OE on cue-induced cocaine seeking (WD1: two-way RM ANOVA, genotype × nosepoke interactions: F_3, 24_ = 4.03, P = 0.02; post hoc, Control virus vs. ANKS1B-OE, P = 0.03; Control virus vs. ANKS1B-OE+FoxO3-OE, P > 0.99; FoxO3-OE vs. ANKS1B-OE, P < 0.01; WD28: two-way RM ANOVA, genotype × nosepoke interactions: F_3, 24_ = 6.23, P < 0.01; post hoc, Control virus vs. ANKS1B-OE, P < 0.01; Control virus vs. ANKS1B-OE+FoxO3-OE, P = 0.58; FoxO3-OE vs. ANKS1B-OE, P < 0.01; n = 7 per group). (I) Timeline of bilateral intra-NAc infusion of the FoxO inhibitor carbenoxolone (150 µM) and behavioral testing. (J) Carbenoxolone infusion did not alter cocaine acquisition during ShA. (K) Inhibition of FoxO3 activity significantly attenuated the escalation of cocaine intake during LgA compared to vehicle control. (L–M) FoxO3 inhibition significantly reduced active nosepokes during cue-induced reinstatement on both WD1 (two-way RM ANOVA, genotype × nosepoke interactions: F_1, 12_ = 4.27, P = 0.06; n = 7 per group) and WD28 (two-way RM ANOVA, genotype effect: F_1, 12_ = 9.74, P < 0.01; post hoc, P < 0.01; n = 7 per group). Data are presented as mean ± SEM. P < 0.05 was considered statistically significant. Statistical significance was determined by two-way repeated-measures (RM) ANOVA followed by Bonferroni’s multiple comparisons test.

## 3. Discussion

This study significantly extends our group’s previous work by delineating a precise molecular mechanism through which ANKS1B, initially identified as a shared genetic risk factor for multiple substance addictions, controls the escalation of cocaine use. While our prior GWAS implicated *ANKS1B* in heroin, methamphetamine, and alcohol dependence, the present findings establish its causal role in cocaine addiction by demonstrating that extended cocaine exposure selectively downregulates ANKS1B expression in the NAc, and that bidirectional manipulation of ANKS1B bidirectionally regulates addiction-like behaviors without affecting natural reward processing. Notably, ANKS1B functions as a pivotal negative regulator of the histone acetyltransferase CBP within the NAc, forming a protein complex that restrains H3K27 acetylation at the FoxO3 promoter. Chronic cocaine exposure disrupts this complex, leading to epigenetic disinhibition of FoxO3 transcription and subsequent maladaptive neuroplasticity. This proposed mechanistic model was present in **Figure 8**, and these results bridge a critical gap between genetic association and functional mechanism, revealing how ANKS1B loss-of-function drives addictive behaviors through a defined CBP-FoxO3 epigenetic pathway. The ANKS1B-CBP-FoxO3 axis thus represents a novel therapeutic target for controlling pathological cocaine escalation.

**Figure 8.**
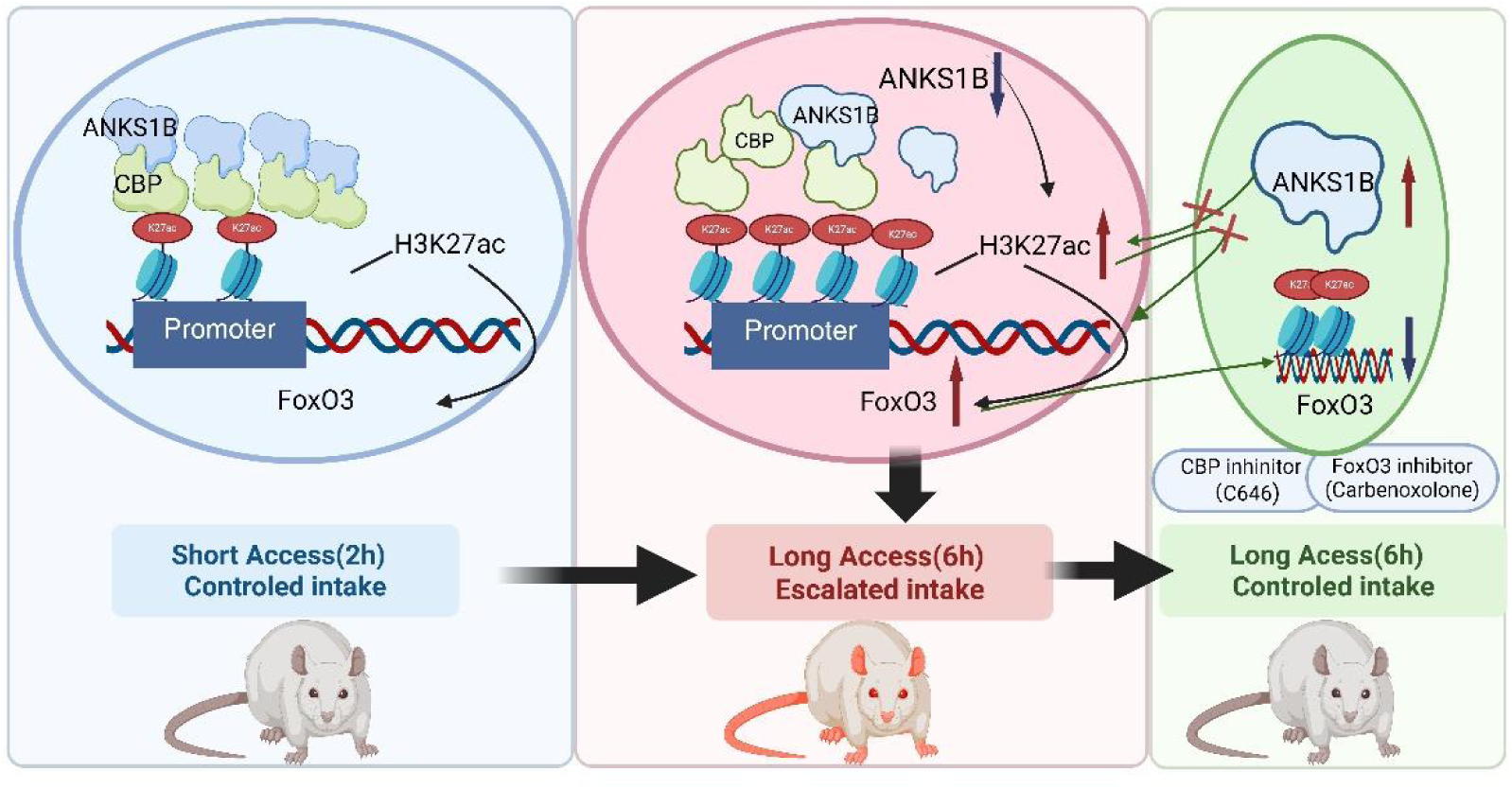
A proposed mechanistic model for ANKS1B/CBP-mediated epigenetic regulation of FoxO3 in the escalation of cocaine seeking. (Left Panel: Short Access/Controlled Intake) Under baseline or short-access cocaine conditions, ANKS1B is bound to CBP at the FoxO3 gene promoter. This complex maintains high levels of the active histone mark H3K27ac, which leads to controlled, low-level expression of Foxo3 and results in a controlled pattern of cocaine intake. (Middle Panel: Long Access/Escalated Intake) During escalated cocaine intake (Long Access, LgA, 6 h/day), ANKS1B expression is downregulated, weakening the ANKS1B–CBP interaction. This disinhibits CBP activity, leading to increased H3K27ac enrichment at the FoxO3 promoter, upregulation of FoxO3 transcription, and facilitation of addiction-related neuroplasticity. (Right Panel: Therapeutic Intervention). Restoring ANKS1B levels (via overexpression) re-establishes the regulatory complex, normalizes H3K27ac and Foxo3 levels, and controls intake. Pharmacological inhibition of downstream targets, such as blocking CBP’s acetyltransferase activity (e.g., with C646) or directly inhibiting Foxo3 activity (e.g., with Carbenoxolone), can bypass the initial deficit in ANKS1B and effectively block the signaling cascade, thereby preventing escalated intake (Image generated: www.biorender.com).

Cocaine self-administration under short- and extended-access conditions induces distinct patterns of cellular and behavioral plasticity (stable low intake and escalated high intake) ^[53,54]^, which was also replicated in our study. Our data showed that NAc ANKS1B expression was downregulated in rats with extended access but not in those with restricted access, which is in line with previous studies showing ANKS1B downregulation across multiple forms of substance abuse, including cocaine ^[55]^, methamphetamine, and heroin ^[17]^. This downregulation is likely attributable to cocaine-induced epigenetic silencing, as chronic cocaine exposure promotes a hypermethylated state in the striatum ^[26, 56–59]^, potentially at ANKS1B regulatory elements ^[60–63]^. Furthermore, cocaine exposure positions *anks1b* loci as binding sites for chromatin remodeling complexes in NAc^[15]^, highlighting its central role as an activity-dependent regulatory molecule. Previous research shows that cocaine use resulted in changes in thin and mushroom spines, which are considered “memory spines” ^[64]^, an effect which is also observed in our current study. Interestingly, ANKS1B-OE mainly influenced cocaine intake-induced changes in thin spines, suggesting that ANKS1B’s role is linked to profound alterations in certain synaptic structural changes. Consistent with the molecular and structural alterations, our behavioral data showed that manipulating ANKS1B expression in the NAc could significantly modulate the escalation of cocaine intake during long-access cocaine sessions, suggesting a critical role of ANKS1B in regulating behavioral and molecular changes associated with excessive cocaine use.

A large body of evidence indicates that as drug use progresses from the controlled ShA stage to an addiction-like LgA state, the underlying neurochemical drivers for drug motivation fundamentally shift, characterized by a significantly enhanced role for glutamatergic signaling and a diminished role for dopamine D1 receptors in NAc ^[35]^. Kasanetz et al. proposed that addiction is linked to a lasting loss of NMDA receptor–dependent LTD in the NAc, whereas this plasticity recovers in non-addicted rats maintaining controlled drug intake ^[65]^. ANKS1B has been shown to play an important role in regulating glutamatergic signaling ^[8]^. In the present study, we found ANKS1B-OE rescued the LgA-induced downregulation of GluN2A, a subunit critical for NMDA receptor function and LTD ^[65–67]^. These results suggest that ANKS1B may serve as a key molecule for maintaining synaptic homeostasis, potentially by regulating glutamatergic signaling, thereby effectively suppressing or constraining cocaine-driven neuroplasticity.

By comparing transcriptional changes in the NAc following short- and long-access cocaine exposure, the predominant effect of cocaine is gene activation, consistent with observations in the NAc ^[68]^. Furthermore, ANKS1B overexpression attenuates the cocaine-induced transcriptional upregulation. The enduring nature of drug addiction, marked by persistent drug-seeking and high relapse rates, points to stable alterations in gene expression programs ^[69–72]^. Histone acetylation, a dynamic and reversible epigenetic modification, stands as a prime candidate for mediating these long-lasting cellular memories ^[69, 73, 74]^. Another intriguing finding from the current study is that ANKS1B-induced changes in NAc acetylation may depend on its binding to CBP. This interaction appears to impede CBP’s ability to acetylate H3K27. It has been suggested that, as a versatile HAT and transcriptional co-activator, CBP can integrate diverse signaling pathways to sculpt gene expression ^[75, 76]^. Accumulating evidence highlights CBP’s established pro-addictive role: its genetic ablation in the NAc significantly attenuates cocaine-induced reward and associated synaptic plasticity, including alterations in dendritic spine morphology ^[76, 77]^. Our findings suggest that prolonged cocaine exposure diminishes the interaction between ANKS1B and the CBP complex, thereby releasing CBP from its inhibitory constraint, resulting in increased CBP-mediated H3K27 acetylation (a critical active enhancer mark) ^[44, 78]^. As a member of the ankyrin repeat–containing scaffold protein family, ANKS1B contains ankyrin repeat domains that function as critical platforms for protein–protein interactions. These domains are known to recruit or modulate epigenetic regulators, including HDACs, thereby directly shaping acetylation homeostasis and downstream transcriptional programs ^[29, 79]^. For instance, ANKRD11, a close homolog of ANKS1B, frequently acts as a transcriptional co-repressor through its interaction with HDACs (e.g. HDAC3), and its loss similarly results in elevated H3K27ac levels ^[80]^. It might be speculated that ANKS1B constrains CBP’s HAT activity through spatial sequestration ^[81]^, conformational alteration ^[82]^, or competitive inhibition ^[83]^. This could be inferred from the roles of other ankyrin-repeat proteins. Collectively, this evidence positions the Ankyrin repeat protein family as versatile organizers of chromatin states, finely tuning gene expression by modulating both HAT (ANKS1B) and HDAC (ANKRD11) activities. Furthermore, large-scale proteomic studies in *Anks1b* haploinsufficient autism models predicted significant alterations in “Sirtuin signaling pathways” ^[9]^, which align with our findings and therefore support the inference that ANKS1B loss perturbs cellular acetylation homeostasis. While further structural and biochemical studies are warranted to dissect these precise modes of action, our data strongly support the conclusion that ANKS1B exerts robust functional inhibition on CBP-mediated H3K27ac.

In the present study, we observed that loss of ANKS1B leads to increased CBP recruitment and H3K27ac at the FoxO3 promoter, whereas ANKS1B overexpression reversed these effects, indicating a direct inhibitory role of ANKS1B in regulating FoxO3. The role of FoxO3 in the central nervous system is context-dependent, encompassing functions in dendritic structure and spine formation, as well as glutamatergic signaling ^[47–49, 84]^. Multiple studies have reported that the FoxO signaling pathway shows significant enrichment across different drugs and various stages of drug use and withdrawal ^[85–87]^. The literature on cocaine-induced FoxO3 signaling is particularly intriguing, revealing a nuanced dual role. Acute cocaine exposure, for instance, can activate an AMPK-FoxO3 pathway that triggers a protective, compensatory autophagy response in the NAc, aimed at counteracting cellular stress ^[88]^. This suggests an initial homeostatic mechanism involving FoxO3. However, during chronic cocaine exposure, FoxO3’s role appears to pivot towards a pro-addictive phenotype. For example, chronic cocaine-induced SIRT1-mediated deacetylation activates FoxO3a, leading to enhanced cocaine reward ^[89]^. Our work provides a crucial, distinct upstream pathway—the ANKS1B-CBP axis—that drives transcriptional upregulation of FoxO3 in long-access cocaine intake. Genome-wide analyses indicate that FoxO transcription factors regulate a wide array of synaptic proteins, including ion channels and receptors, underscoring their capacity to remodel synaptic composition ^[90, 91]^. Building on this framework, our data suggest that ANKS1B downregulation–driven FoxO3 induction may contribute to reduced *grin2a* (NR2A) expression and dendritic spine remodeling, processes further reinforced by the known reciprocal regulatory interactions between NMDAR signaling and FoxO activity. Nevertheless, more studies are needed to directly test whether FoxO3 is the causal mediator of these synaptic alterations.

In summary, we demonstrated a novel regulatory axis in which ANKS1B functions as a crucial inhibitory scaffold for CBP, controlling H3K27ac levels and the expression of the pro-addictive transcription factor FoxO3. Long-term cocaine-induced downregulation of ANKS1B removes this brake, leading to epigenetic and transcriptional disinhibition that drives pathological plasticity. This work reframes addiction as a disorder involving not only aberrant activation of signaling pathways but also the failure of endogenous inhibitory mechanisms that maintain epigenetic homeostasis. Our data suggest that manipulating the ANKS1B-CBP-FoxO3 axis representing a promising therapeutic target for excessive cocaine use.

## 4. Experimental Section

### Experimental Animals and Housing Conditions

Male Sprague-Dawley rats, weighing between 260 and 280 g, were procured from Beijing Vital River Laboratory Animal Technology Co., Ltd. Before the experiments, the rats were group-housed with five animals per cage. The animal facility maintained a controlled environment with a temperature of 23–27°C and humidity of 50±5%. A reverse 12-hour light/dark cycle was implemented, and the animals were provided with unrestricted access to standard chow and water. All experimental protocols were conducted in strict accordance with the Guide for the Care and Use of Laboratory Animals from the National Institutes of Health and received approval from the Biomedical Ethics Committee for Animal Use and Protection at Peking University Health Science Center (Animal Protocol Approval Number: DLASBE0353).

### Surgery

In the self-administration paradigm, rats were anesthetized with isoflurane, and a silastic catheter was implanted into the right jugular vein with the distal end positioned at the entrance of the right atrium, following established procedures ^[92]^. To ensure catheter patency and prevent infection, daily flushing was performed with heparin sodium (4 mg/ml in 0.9% saline) and benzylpenicillin sodium (200,000 U/ml in 0.9% saline).

For intracranial drug or viral delivery, permanent 23-gauge guide cannulas were stereotaxically implanted bilaterally, positioned 1 mm above either the NAc. The stereotaxic coordinates used were as follows: AP: −1.1 mm relative to bregma, ML: ±1.2 mm, DV: −7.4 mm. Following surgery, animals were singly housed and given 3–5 days to recover before the onset of behavioral testing. Cannula locations were confirmed by immunofluorescence, and rats with incorrect placements were excluded from subsequent analyses.

### Drug and virus injection procedures

Carbamazepine (CBZ), c646 were purchased from Sigma Aldrich (St. Louis, MO, USA). Each drug (0.5 μl/side) was delivered into the NAc via chronically implanted cannulae, with a Hamilton 10 μl syringe pump at a rate of 3 μl/10 min. The final concentration that was injected into the Acb for the individual drugs was CBZ (150 μmol) ^[93]^, c646 (500 ng/side) ^[94]^. Recombinant adeno-associated virus (rAAV) vectors that contained shRNA that targeted ANKS1B (rAAV-hSyn-sh*Anks1b*-mCherry), overexpression virus (AAV-CMV-*Anks1b*-FLAG-WPRE), control virus (AAV-Flag or AAV-hSyn-Scramble-mCherry) were commercially purchased (5.00 × 10^12^ viral genomes/ml; BrainVTA Co. Ltd., Wuhan, China). Lentiviral overexpression of FoxO3 was performed using a GV492 vector (GeneChem, Shanghai, China) encoding the full-length rat FoxO3 under the control of the ubiquitin (Ubc) promoter. The construct contained a 3×FLAG tag and a GFP reporter linked via an IRES element (Ubc-MCS-3FLAG-CBh-GFP-IRES-puromycin). Recombinant lentivirus (LV-Foxo3) was produced by GeneChem with a final titer of 2.20 × 10^8 TU/mL. Lentiviral overexpression of ANKS1B was performed using a KV927 vector (GeneChem, Shanghai, China) encoding the full-length rat ANKS1B (NM_001414940.1) under the control of the hSyn promoter. The construct included an EGFP reporter and a 3×FLAG tag for visualization and detection (hSyn-MCS-EGFP-3FLAG-SV40-puromycin). Recombinant lentivirus (LV-Anks1b) was produced and purified by GeneChem with a final titer of 3.30 × 10^9 TU/mL. An EGFP-expressing lentiviral vector was used as the control. The virus was injected with a volume of 500 nl per side into the NAc. The injections lasted over 3 min, and the needles were kept for an additional 10 min to allow for the drug or virus to be completely infused.

### Cocaine Self-Administration Paradigm

Rats weighing 280–300g at the start of the study underwent surgery for the implantation of a chronic indwelling jugular catheter under anesthesia with a ketamine/xylazine mixture (75 and 5 mg·kg⁻¹, i.p.), following previously established procedures ^[30]^. A recovery period of at least one week was allowed post-surgery. To maintain catheter patency and prevent infection, catheters were flushed daily for one week with a 0.2 ml solution of enrofloxacin (4 mg·ml⁻¹) in heparinized saline (50 IU·ml⁻¹).

The self-administration training began with an acquisition phase where rats were trained on a short-access schedule (2 h per session) to lever-press for intravenous cocaine infusions (0.5 mg·kg⁻¹ per infusion). Responses on the active lever were reinforced on a fixed-ratio (FR) schedule, followed by a 30-second time-out period. Each infusion was paired with a 5-second illumination of a stimulus light located above the active lever, and the main chamber light was turned off during the time-out. Sessions concluded after either 2 hours had elapsed or 40 infusions had been earned. Over seven days, the response requirement was progressively increased from FR1 to FR5. Rats that achieved more than 10 infusions in a single session were advanced to the next FR schedule. By day 7, all rats were on a fixed-ratio (FR) 5 schedules. For the extended-access experiments, rats underwent a 10-day maintenance period on a long-access schedule (6 h per session).

Cue-induced cocaine-seeking was assessed either 1 or 28 days after the 10-day self-administration period. Rats were housed in their home cages during the abstinence phase with no extinction training. For the seeking test (2 h duration), active lever presses on an FR5 schedule resulted in the presentation of the light cue as in the training phase, but no cocaine was delivered.

### Immunofluorescence Staining

Immunofluorescence was performed on 25-µm-thick coronal brain sections as previously described ^[95, 96]^. The following primary antibodies were used: rabbit anti-ANKS1B (1:500, Proteintech), mouse anti-GFAP (1:500, Abcam), mouse anti-NeuN (1:500, CST), and mouse anti-CBP (1:100, Santa Cruz). Secondary detection was achieved using: Alexa Fluor 594-labeled goat anti-rabbit antibody and Alexa Fluor 488-labeled goat anti-mouse antibody (both 1:1000 dilution, Life Technologies).

### Western Blotting Analysis

Western blotting was conducted following established protocols ^[95]^. The primary antibodies included: rabbit anti-ANKS1B (1:500, Proteintech), mouse anti-H3 (1:2000, CST), mouse anti-H3K9ac (1:11000, CST), mouse anti-H3K14ac (1:11000, CST), mouse anti-H3K27ac (1:1000, CST), mouse anti-H3K56ac (1:1000, CST), mouse anti-H2bK12ac (1:1000, beyotime), mouse anti-H2b (1:1000, beyotime). A horseradish peroxidase-conjugated secondary antibody (goat anti-rabbit and goat anti-mouse IgG, 1:2000, Zsbio) was used for detection.

### Co-immunoprecipitation (Co-IP)

Tissue samples were homogenized in an IP lysis buffer (50 mM Tris-HCl, pH 7.4, 150 mM NaCl, 1 mM EDTA, 0.3% NP-40) containing a protease inhibitor cocktail, followed by sonication and centrifugation. The resulting protein supernatant was incubated for 6 hours at 4°C with 2 µg of the CBP (Santa Cruz) and 20 µL of protein A/G agarose beads (Santa Cruz). The immunoprecipitates were then washed, eluted from the beads, and subsequently analyzed by Western blotting.

### Quantitative Real-Time PCR (qPCR)

Total RNA was extracted from isolated NAcc tissue samples using TRIzol reagent (Invitrogen). The RNA was then reverse transcribed into cDNA using the HiScript II 1st Strand cDNA Synthesis Kit (Vazyme Biotech). qPCR was performed using SYBR Green SuperMix (TOYOBO) on a QuantStudio 5 Real-Time PCR System (Applied Biosystems). Each 20 µl reaction contained 10 µl of 2× master mix, 2 µl of cDNA, 1 µl of each primer, and 6 µl of distilled water. The thermal cycling parameters were: an initial denaturation at 95°C for 10 minutes, followed by 40 cycles of 95°C for 30 seconds and 60°C for 1 minute. All reactions were run in duplicate, and a melting curve analysis was performed. Gene expression was quantified using the ΔΔCt method, with Actin serving as the reference gene. The primer sequences for Foxo1, Foxo3, and Foxo6 were either designed using Primer Premier 5.0 or obtained from previous publications (listed in Supplementary Table 1).

### Chromatin Immunoprecipitation (ChIP)

Cross-linking was performed by adding 37% formaldehyde directly to the culture medium to a final concentration of 1% and incubating for 10 minutes at 37°C. The reaction was quenched by adding glycine to a final concentration of 125 mM. The ChIP procedure was carried out using a ChIP Assay Kit (Cell Signaling Technology) according to the manufacturer’s protocol. The primary antibodies included CBP (1:50, CST) and H3K27ac (1:50, CST). The extracted DNA was then analyzed by ChIP-qPCR and ChIP-Sequencing using specific primers for the FOXOs genome (listed in Supplementary Table 2).

### RNAscope in situ Hybridization

In order to explore the co-expression of *Anks1b* and intermediate spiny neurons, RNA in situ hybridization was performed on 25-µm-thick coronal sections as previously described ^[97]^. The procedure utilized probes targeting the mRNA transcripts for Drd1, Drd2, and *Anks1b* (1:25, Advanced Cell Diagnostics). Drd1 and Drd2 were visualized with Opal 570 and Anks1b with Opal 690 (1:100, Akoya Biosciences).

### Behavioral Assays

All behavioral tests were conducted during the dark cycle, adapted from previous reports with minor modifications ^[98]^.

### Open Field Test (OFT)

To assess general locomotor activity and anxiety-like behavior, rats were placed in a corner of a wooden open-field apparatus (100 × 100 × 40 cm). The arena was divided into 16 equal squares, with the central four squares defined as the “center zone.” The total distance traveled and the time spent in the center zone during a 5-minute session were recorded and analyzed using EthoVision XT 10 software (Noldus).

### Elevated Plus Maze (EPM) Test

Anxiety-like behavior was further evaluated using the EPM, which was elevated 70 cm from the floor. The maze consisted of two open arms (50 × 12 cm) and two closed arms (50 × 12 cm with 30 cm high walls) arranged in a plus shape. Each rat was placed in the central platform facing an open arm and allowed to explore freely for 5 minutes. The number of entries into and the time spent in the open arms were automatically recorded using EthoVision XT 10 software.

### Novel Object Recognition (NOR) Test

The NOR test was adapted from a previously published protocol and conducted in two identical black open-field boxes (85 × 85 × 55 cm). The objects used were made of glass or plastic and were too heavy for the rats to move. Habituation consisted of two 10-minute sessions on consecutive days, where rats explored the empty box. The test was conducted on the third and fourth days. Each day involved a 3-minute training session with two identical objects (A/A), followed by a 3-minute test session either 1 or 24 hours later with one familiar and one novel object (A/B). The time spent exploring each object was recorded. A discrimination index was calculated for the A/B sessions as: (Time exploring novel object - Time exploring familiar object) / (Total exploration time).

### RNA-seq

Post-behavioral assessments: Tissue from the NAc was collected for RNA extraction, which was then used for RNA analysis (APExBIO). GO and KEGG analyses of differentially expressed genes were conducted using the online tool DAVID database (https://david.ncifcrf.gov/home.jsp) and obtained in SCV format for GO and KEGG enrichment. The KEGG pathways with p < 0.05 were extracted and visualized using the online tool Weishengxin (https://www.bioinformatics.com.cn/). Within each comparison group, the results were considered significant at the following thresholds: p value ≤ 0.05 and absolute value of |log2(fold change) | ≥ 1.2 ^[99]^.

### Golgi-Cox Staining

Animals were then deeply anesthetized with sodium pentobarbital (100 mg/kg, i.p.) and perfused transcardially with 0.9% saline. Brains were removed and processed using the Golgi-Cox method as described by Glaser and Van der Loos ^[100]^. In brief, tissue was immersed in Golgi-Cox solution at room temperature for 14 days, transferred to 30% sucrose, and subsequently sectioned at 100 μm thickness on a vibratome. Coronal slices were mounted on gelatin-coated slides, stained following established procedures ^[101]^, coverslipped, and air-dried before quantitative assessment. From each group, 3–5 neurons per rat were analyzed, with 5–6 animals contributing to the dataset; mean values per animal were then used for statistical comparisons. Images were acquired using an Olympus BX53 microscope equipped with a 100× oil-immersion objective. Dendritic length was quantified with NIH ImageJ software, and dendritic spine counts were performed independently by two blinded experimenters. Spine density was expressed as the average number of spines per 10 μm of dendritic segment.

### Bioinformatics analyses

Protein-protein complex structures were predicted using AlphaFold3 (https://alphafoldserver.com/). Amino acid sequences of the target proteins were obtained from UniProt (https://www.uniprot.org/) and used as input. Multiple sequence alignments (MSAs) were automatically generated by the AF3 pipeline. Structural predictions were sampled from multiple random seeds, each producing candidate models through iterative diffusion and denoising steps. Following the recommended inference protocol, multiple models were generated and ranked based on predicted template modeling (pTM) and interface predicted TM-score (ipTM), and the top-ranked structure was selected for further structural analysis (listed in Supplementary Table 3). Interface reliability was evaluated based on ipTM in combination with Predicted Aligned Error (PAE) and predicted Local Distance Difference Test (pLDDT) scores, with higher ipTM, lower inter-chain PAE, and consistently high pLDDT indicating more reliable interactions. Visualization of the final models was performed using PyMOL (The PyMOL Molecular Graphics System, Version 3.1.6.1, Schrödinger, LLC).

## Statistical analysis

All statistical analyses were performed using GraphPad Prism software (version 8.0). Data are presented as mean ± SEM. Depending on the experimental design, one-way or two-way analysis of variance (ANOVA) was performed with appropriate between-subject or within-subject factors. For between-subject factors, post hoc comparisons were conducted using Tukey’s HSD test. For within-subject (repeated-measures) factors, post hoc comparisons were carried out using Bonferroni-corrected pairwise t-tests or Dunnett’s test, as appropriate. For comparisons between two groups, unpaired t-tests were applied when the data met assumptions of normality and homogeneity of variance. Statistical significance was defined as p < 0.05.

## Author Contributions

L.L., J.S., J.L., and Y.S. conceived the project. L.Y., X.C., C.P., Z.L., J.D., W. D., S.G., and L. Lv. performed the experiments. L.Y., J.L., and Y.S. contributed to data analysis and interpretation. S.M. contributed to funding support. Y.X. and Y.H. provided help with the experiments and experimental design. L.Y., J.L., and Y.S. wrote and revised the manuscript, with input from all authors. All authors discussed the results and approved the final version of the manuscript.

## Acknowledgements

This study was supported by funding from the National Natural Science Foundation of China (No. 82171488, 82471515 and 82271533), and STI2030-Major Projects (No. 2021ZD0201900). The authors acknowledge BioRender for providing the scientific image (License n. ZC29OF6QUP).

## Data Availability Statement

The transcriptome dataset generated in this study has been deposited in the National Center for Biotechnology Information (NCBI) database as a BioProject (accession no. PRJNA1338099). The datasets used and/or analyzed during the current study are available from the corresponding author on reasonable request.

## Supporting Information

**Figure S1.**
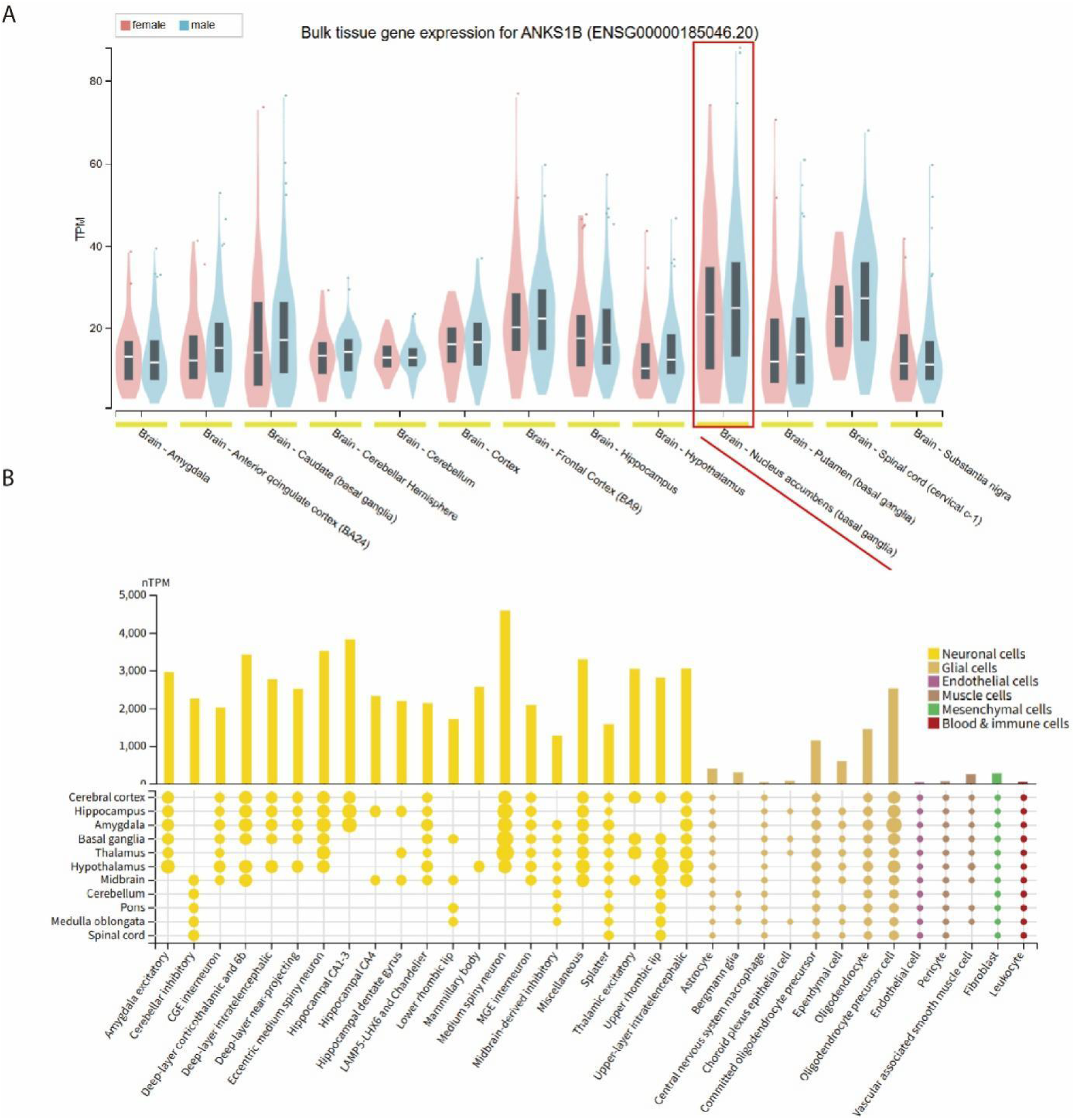
Expression profile of *ANKS1B* in the brain. **(A)** Violin plot showing the gene expression level (TPM: Transcripts Per Million) of *ANKS1B* in different regions of the humanbrain from the GTEx database bulk RNA-sequencing data. The data indicate that *ANKS1B* is expressed across multiple brain regions, including the Nucleus Accumbens (NAc, highlighted by the red box). **(B)** Single-cell transcriptomics data showing *ANKS1B* expression (nTPM: normalized Transcripts Per Million) in various brain cell types. The bar plot on top shows that *ANKS1B* is highly expressed predominantly in neuronal cells (yellow). The dot plot below further illustrates its distribution among specific neuronal subtypes.

**Figure S2.**
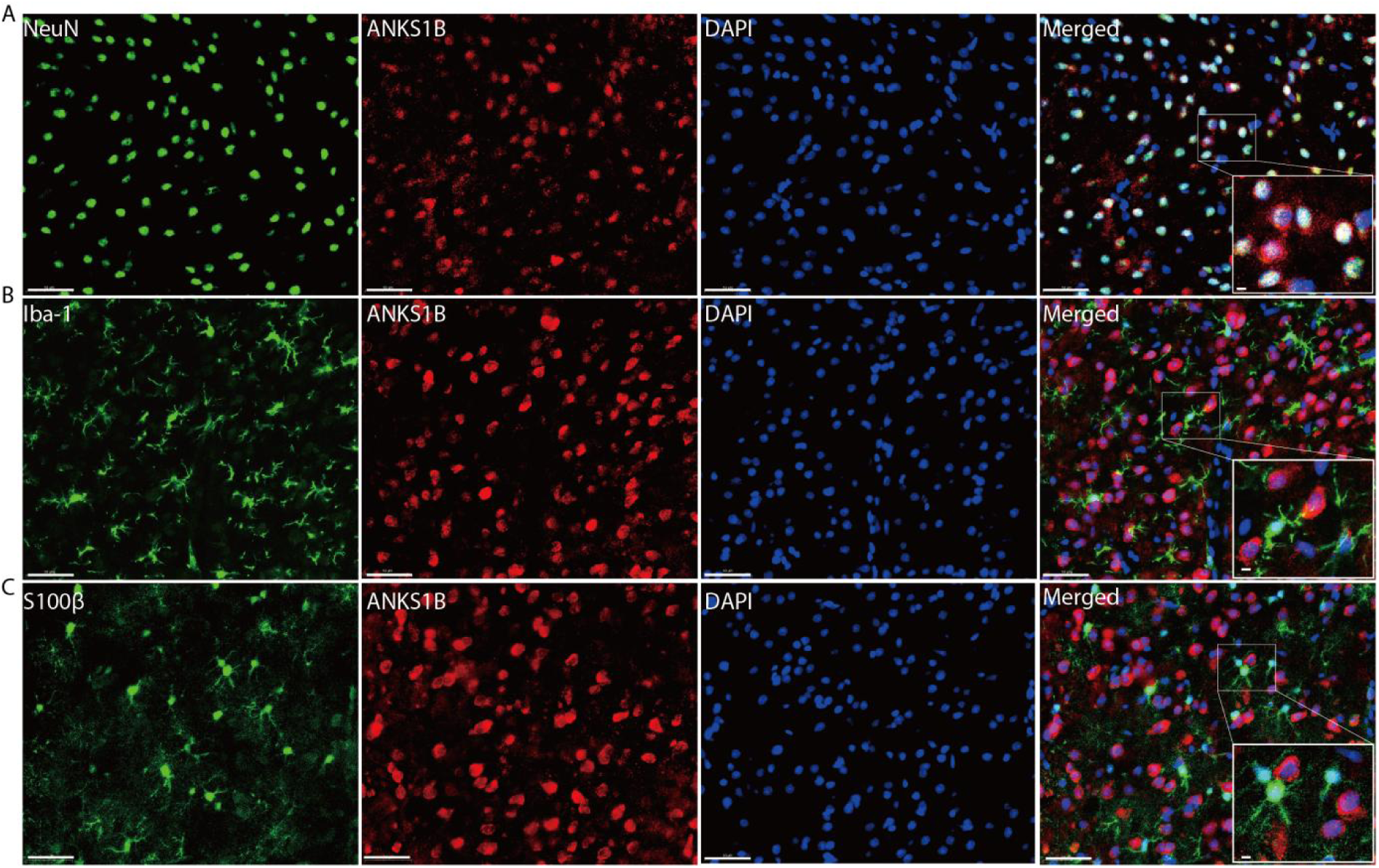
ANKS1B is predominantly expressed in neurons within the NAc. **(A)** Immunofluorescence co-staining shows extensive co-localization of ANKS1B (red) with the neuronal marker NeuN (green), indicating ANKS1B expression in neurons. **(B)** Co-staining of ANKS1B (green) with the microglial marker Iba-1 (green) shows no co-localization. **(C)** Co-staining of ANKS1B (red) with the astrocyte marker S100β (green) shows little to no co-localization. All scale bars, 50 µm and 5 µm.

**Figure S3.**
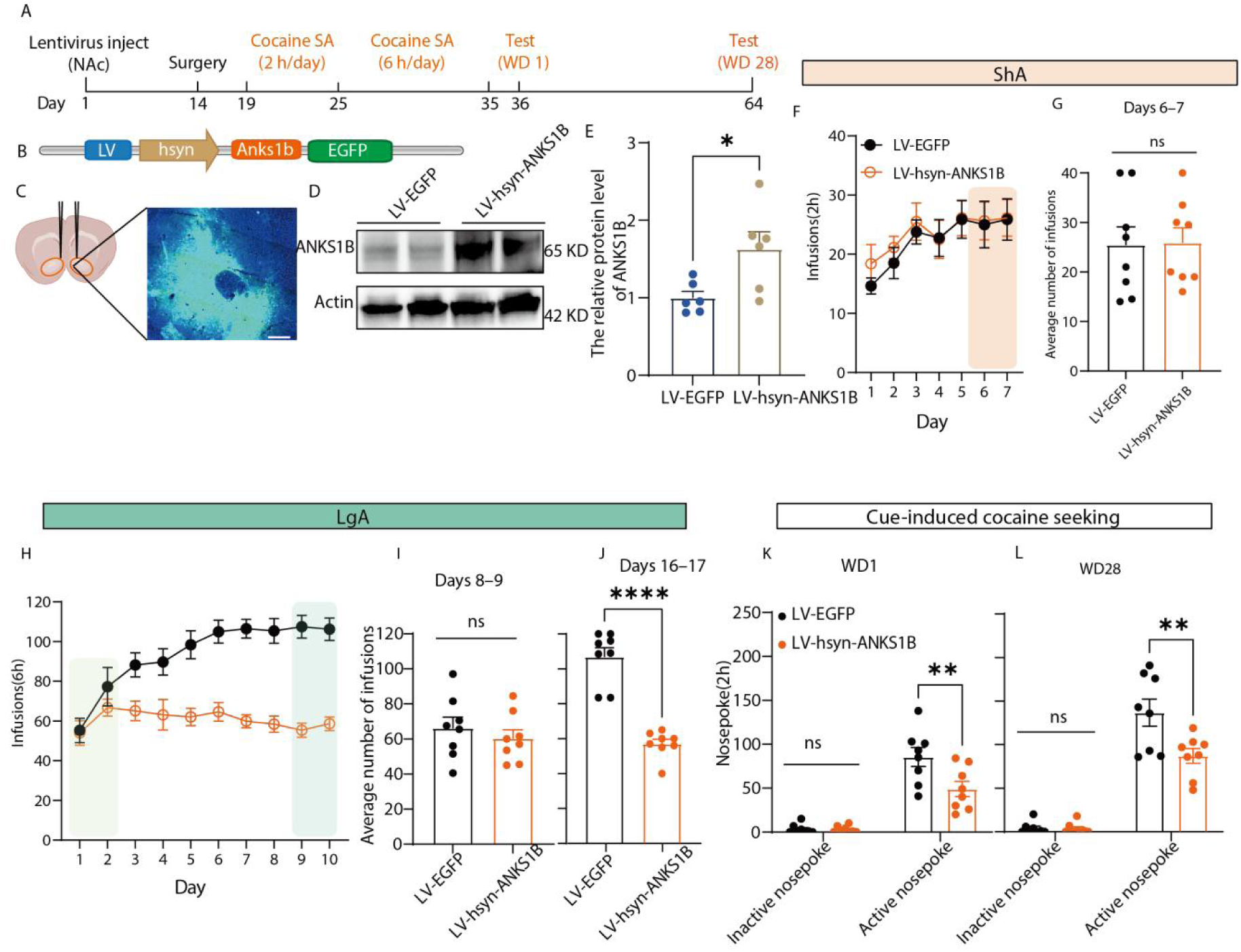
ANKS1B overexpression in the NAc suppresses escalation of cocaine self-administration and cue-induced seeking. **(A)** Experimental timeline showing lentivirus injection, cocaine self-administration, and WD testing. **(B)** Representative image showing LV-hsyn-ANKS1B-EGFP expression in the NAc. Scale bars, 100 μm. **(C)** Western blot validation confirming successful ANKS1B overexpression in the NAc compared with the control group (unpaired t-test, t_10_ = 2.66, P = 0.02, n = 6 per group). **(D)** ANKS1B overexpression did not alter cocaine acquisition during the ShA phase (n = 8 per group). **(E)** Comparison of cocaine infusions during the last 2 days of ShA (days 6–7) revealed no difference between groups (unpaired t-test, t_14_ = 0.09, P = 0.93). **(F)** ANKS1B overexpression prevented escalation of cocaine intake during the LgA phase. **(G)** Comparison of infusions on the first 2 days (days 8–9; unpaired t-test, t_14_ = 0.74, P = 0.47) and the last 2 days (days 16–17; t_14_ = 8.35, P < 0.01) of LgA. **(H and I)** Overexpression of ANKS1B significantly reduced cue-induced cocaine seeking (Active nosepoke) on both WD1 and WD28. (WD1: two-way RM ANOVA, genotype × nosepoke interactions: F_1, 14_ = 6.60, P = 0.02; post hoc, P = 0.01; WD28: two-way RM ANOVA, genotype × nosepoke interactions: F_1, 14_ = 6.64, P = 0.02; post hoc, P < 0.01; n = 8 per group). An EGFP-expressing lentiviral vector (LV-EGFP) was used as the control. Data are presented as mean ± SEM. P < 0.05 was considered statistically significant. Statistical significance was determined by two-way repeated-measures (RM) ANOVA followed by Bonferroni’s multiple comparisons test.

**Figure S4.**
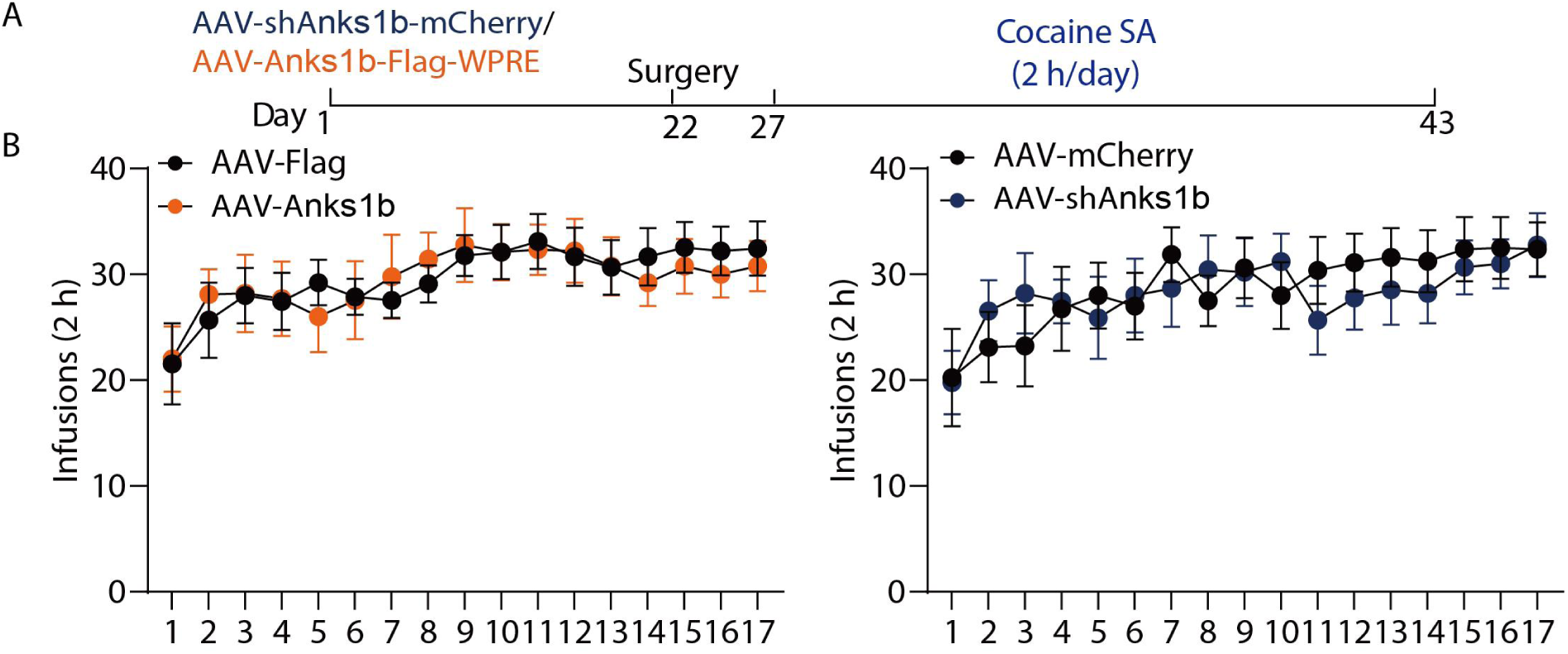
Manipulation of ANKS1B levels in the NAc does not affect Short-access cocaine acquisition of short-access cocaine self-administration. **(A)** Experimental timeline showing a 17-day acquisition period of cocaine self-administration (SA) under a short-access (2h/day) schedule. **(B)** Daily cocaine infusions. The left panel shows that overexpression of ANKS1B did not alter drug intake over the 17-day acquisition period compared to the control virus group. The right panel shows that knockdown of ANKS1B also did not affect drug intake (Two-way ANOVA, p > 0.05). n = 7 per group. Data are presented as mean ± SEM.

**Figure S5.**
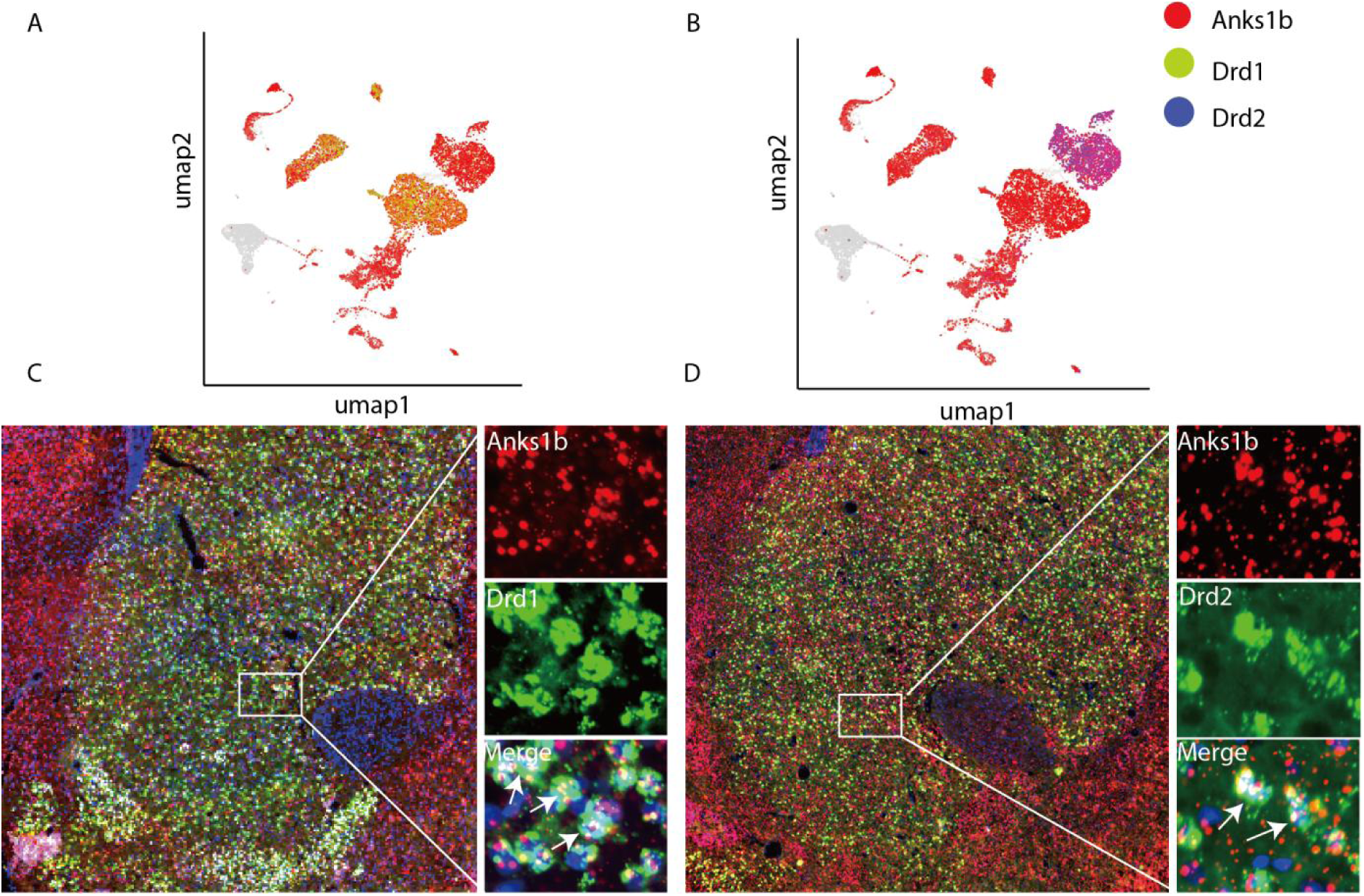
*Anks1b* is expressed in both D1- and D2-receptor-expressing medium spiny neurons (MSNs) in the NAc. **(A–B)** UMAP (Uniform Manifold Approximation and Projection) plots of single-cell RNA-sequencing data from the NAc. A) shows the expression distribution of the *Anks1b* gene (red indicates high expression). B) shows the expression distribution of dopamine D1 receptor (*Drd1*, green) and D2 receptor (*Drd2*, blue). Comparison of the plots indicates that *Anks1b* is expressed in both D1-MSNs and D2-MSNs. **(C–D)** Fluorescent in situ hybridization (FISH) images confirming the co-localization of *Anks1b* mRNA (red) with *Drd1* mRNA (green) in NAc neurons (yellow signal in Merge panels, indicated by arrows).

**Figure S6.**
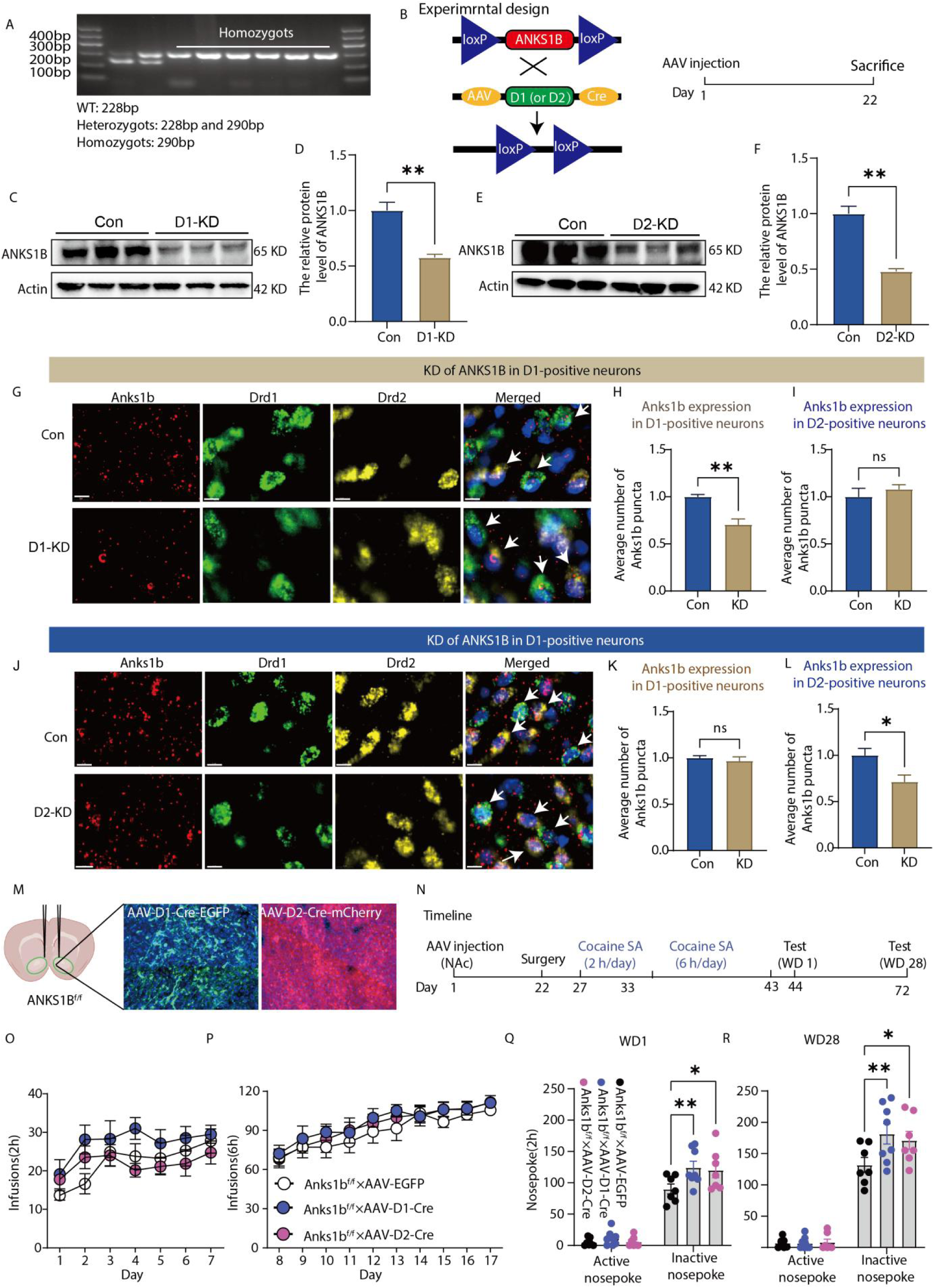
Cell-type–specific deletion of ANKS1B in NAc D1- or D2-MSNs regulates cocaine seeking. **(A)** RPCR genotyping of Anks1b^f/f^ rats showing WT (228 bp), heterozygous (228/290 bp), and homozygous (290 bp) bands. **(B)** Schematic workflow of the experimental design. AAV-D1-Cre or AAV-D2-Cre was injected into the NAc of Anks1b^f/f^ rat to induce site-specific recombination of loxP sites. Tissues were harvested for analysis 21 days post-injection. **(C-D)** Western blot validation and quantification of ANKS1B protein levels in the NAc after D1-KD (unpaired t-test, t_4_ = 5.30, P = 0.01, n = 3 per group). **(E–F)** Western blot validation and quantification after D2-KD (unpaired t-test, t_4_ = 7.37, P < 0.01, n = 3 per group). Representative RNAscope images of Anks1b mRNA (red) with Drd1 (green) and Drd2 (yellow) labeling in the NAc. In Anks1b ^f/f^ × AAV-D1-Cre animals (D1-MSN-specific ANKS1B KD), Anks1b puncta are selectively reduced in Drd1-positive neurons but not in Drd2-positive neurons. Arrows indicate colocalized puncta. All scale bars, 10 µm. **(G)** Representative RNAscope images of Anks1b mRNA (red) with Drd1 (green) and Drd2 (yellow) labeling in the NAc. In Anks1b^f/f^ × AAV-D1-Cre animals (D1-MSN-specific ANKS1B KD), Anks1b puncta are selectively reduced in Drd1-positive neurons but not in Drd2-positive neurons. Arrows indicate colocalized puncta. All scale bars, 10 µm. H, I) Quantification of Anks1b puncta in D1- and D2-MSNs following D1-KD. Anks1b signal is significantly reduced in D1-MSNs but unchanged in D2-MSNs (unpaired t-test, t_6_ = 4.62, P < 0.01; t_6_ = 0.80, P = 0.46; n = 4 per group). **(H–I)** Quantification of Anks1b puncta in D1- and D2-MSNs following D1-KD. Anks1b signal is significantly reduced in D1-MSNs but unchanged in D2-MSNs (unpaired t-test, t_6_ = 4.62, P < 0.01; t_6_ = 0.80, P = 0.46; n = 4 per group). **(J)** Representative RNAscope images of Anks1b mRNA (red) with Drd1 (green) and Drd2 (yellow) labeling in the NAc. In Anks1b^f/f^ × AAV-D2-Cre animals (D2-MSN-specific ANKS1B KD), Anks1b puncta are selectively reduced in Drd2-positive neurons but not in Drd1-positive neurons. Arrows indicate colocalized puncta. All scale bars, 10 µm. **(K–L)** Quantification of Anks1b puncta following D2-KD. Anks1b expression is selectively reduced in D2-MSNs, with no effect in D1-MSNs (unpaired t-test, t6 = 0.63, P = 0.55; t6 = 2.74 P = 0.03, n = 4 per group). **(M)** Representative images confirming successful infection of NAc neurons. Left: schematic illustration of injection sites in the NAc. Middle: AAV-D1-Cre-EGFP expression (green) in NAc. Right: AAV-D2-Cre-mCherry expression (red) in NAc. **(N)** Experimental timeline. After viral injection and recovery, rats underwent short-access cocaine self-administration (SA; 2 h/day) for 7 days, followed by long-access cocaine SA (6 h/day) for 10 days. Drug-seeking behavior was tested on WD1 and WD28 under extinction conditions. **(O)** Cocaine intake during 2 h/day SA sessions. Deletion of ANKS1B in either D1- or D2-MSNs did not alter cocaine intake relative to controls. **(P)** Cocaine intake during 6 h/day SA sessions (long access). All groups exhibited escalation of intake across training days, with no significant differences between groups. **(Q)** Cocaine seeking at WD1. A two-way RM analysis revealed a significant main effect of genotype (F_2, 19_ = 5.48, P = 0.01), with no significant genotype × lever interaction (F_2, 19_ = 2.27, P = 0.12). Post hoc Dunnett’s multiple-comparisons test showed that both Anks1b^f/f^ × D1-Cre and Anks1b^f/f^ × D2-Cre groups exhibited significantly increased active nosepokes compared with controls (P = 0.01 and P = 0.02, respectively), whereas no differences were observed in inactive nosepokes. **(R)** Cocaine seeking at WD28. A two-way RM analysis revealed a significant lever × genotype interaction (F_2, 19_ = 4.58, P = 0.02). Post hoc Dunnett’s multiple-comparisons test showed that both Anks1b^f/f^ × D1-Cre and Anks1b^f/f^ ×D2-Cre groups exhibited significantly increased active nosepokes compared with controls (p < 0.01 and p = 0.03, respectively), whereas no differences were observed in inactive nosepokes. n = 7–8 per group. Data are presented as mean ± SEM. P < 0.05 was considered statistically significant. Statistical analysis was performed using two-way ANOVA followed by Dunnett’s post hoc multiple-comparisons test.

**Figure S7.**
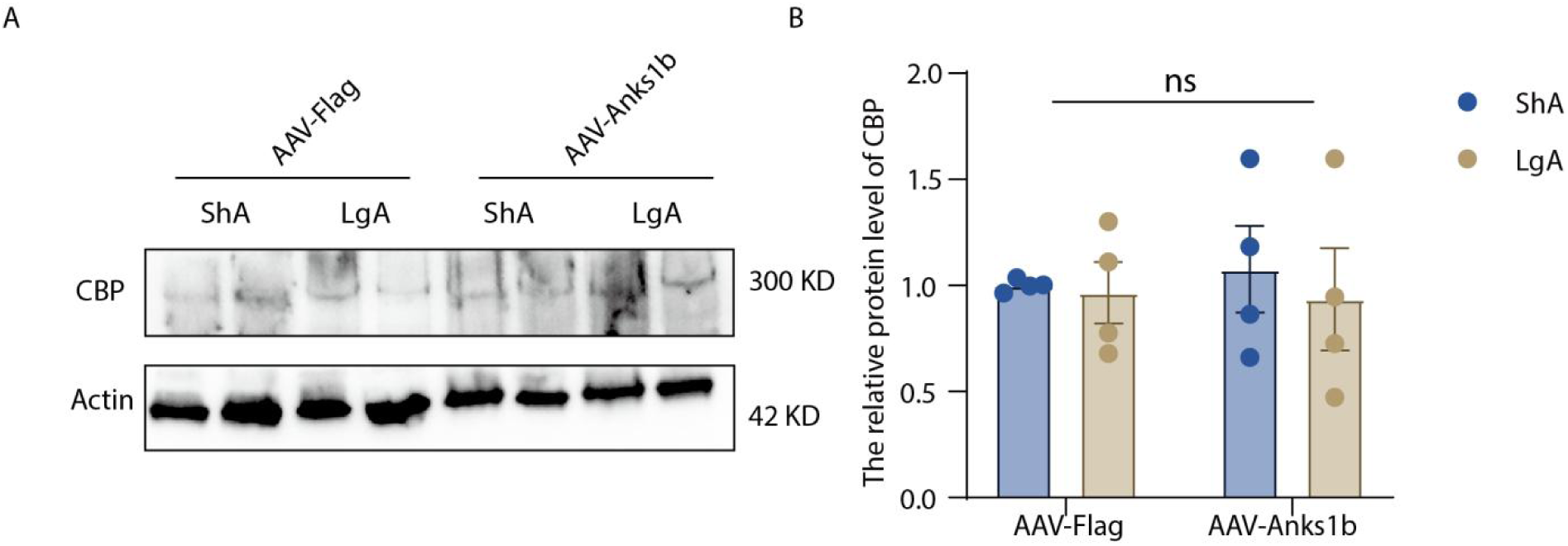
ANKS1B overexpression does not alter total CBP protein levels in the NAc following cocaine self-administration. **(A–B)**.Representative western blot images and quantitative analyses indicate that neither extended cocaine self-administration nor ANKS1B overexpression alters CBP protein levels in the nucleus accumbens. (two-way ANOVA, genotype × access interaction: F_1, 12_ = 0.09, P = 0.76; n = 4 per group). Data are presented as mean ± SEM. P < 0.05 was considered statistically significant. Statistical significance was determined by two-way ANOVA followed by Tukey’s post hoc test.

**Figure S8.**
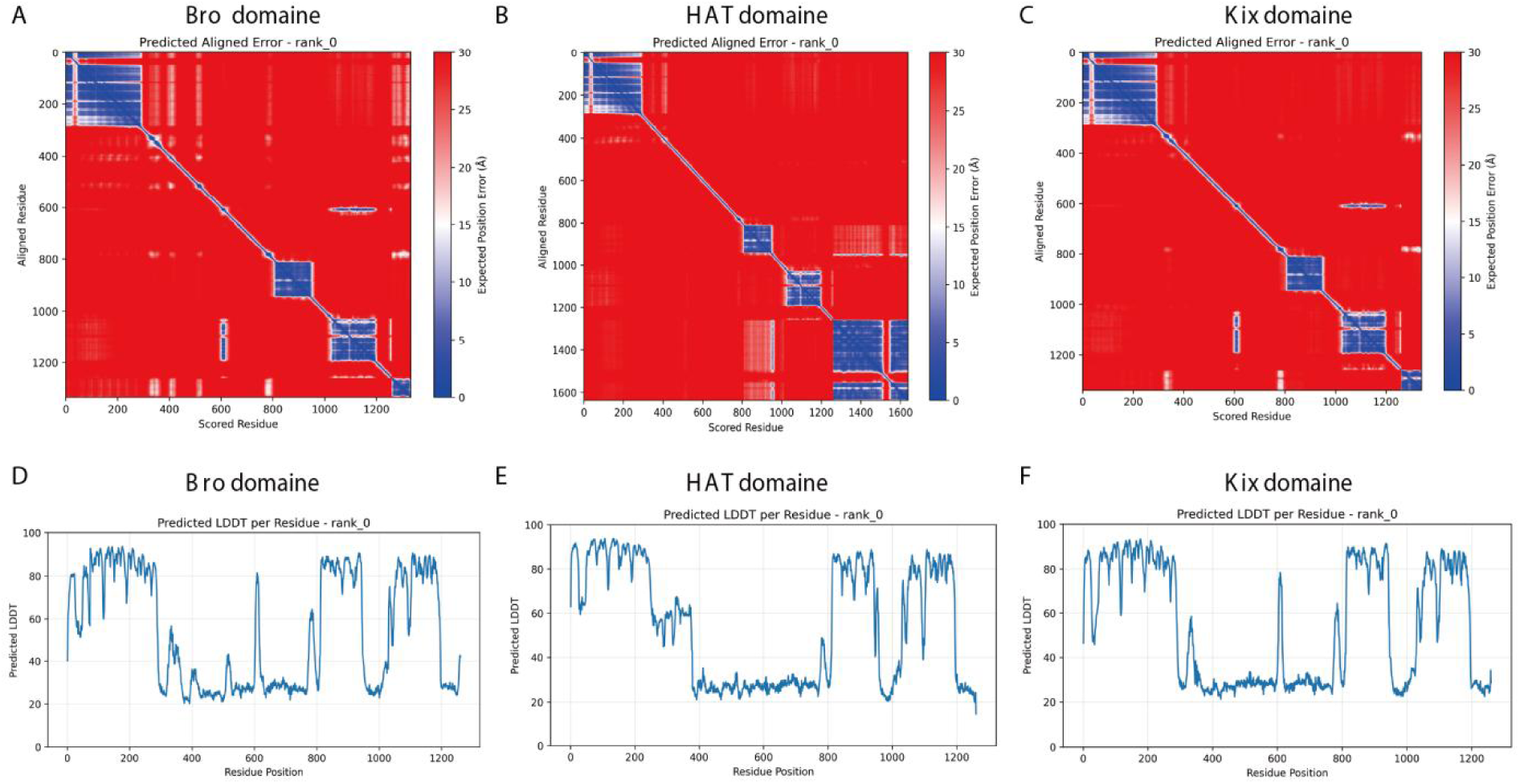
Structural prediction confidence of ANKS1B-CBP domain interactions based on AlphaFold analysis. (A–C) Predicted aligned error (PAE) plots for ANKS1B in complex with CBP domains, including the Bro domain (A), HAT domain (B), and KIX domain (C). Lower PAE values (blue) indicate higher confidence in the relative positioning of residues, whereas higher values (red) indicate lower confidence. **(D–F)** Predicted local distance difference test (pLDDT) scores per residue for the corresponding complexes shown in (A–C), including the Bro domain (D), HAT domain (E), and KIX domain (F). Higher pLDDT scores indicate greater confidence in local structural predictions.

**Figure S9.**
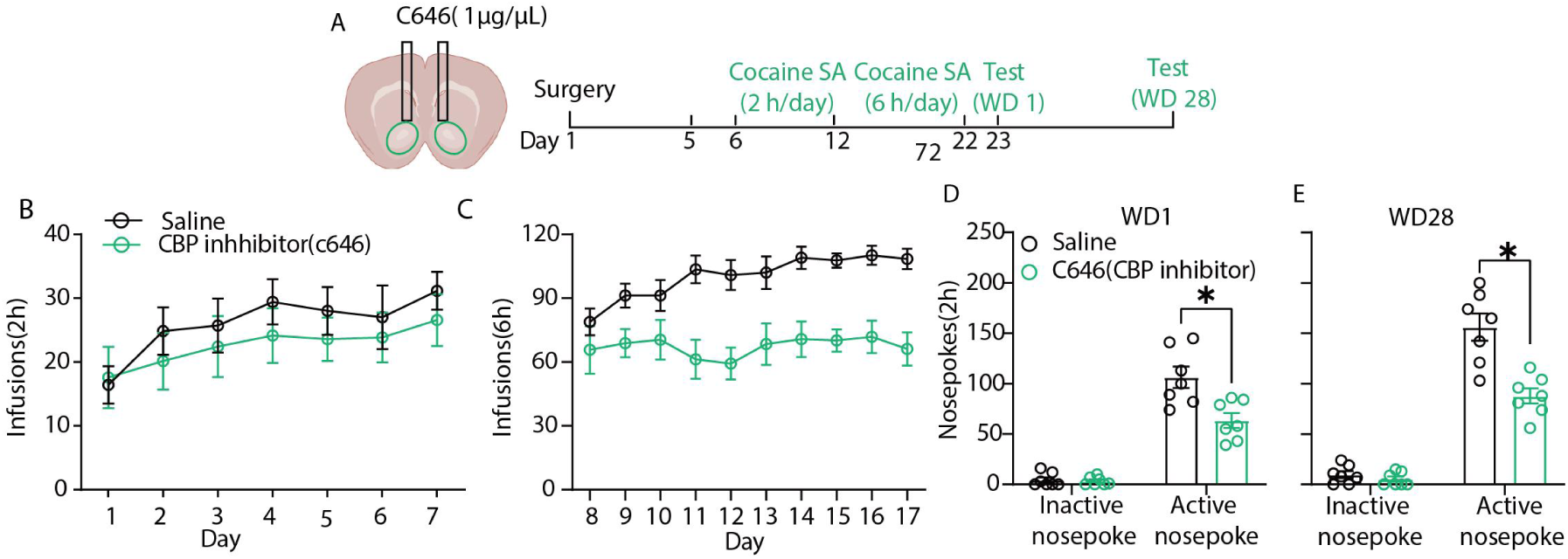
Pharmacological inhibition of CBP in the NAc promotes escalated cocaine intake and seeking. **(A)** Schematic of the experimental design, involving direct infusion of the CBP inhibitor C646 into the bilateral NAc. **(B)** C646 had no significant effect on cocaine intake during the short-access (2h/day) phase. **(C)** During the long-access (6h/day) phase, C646 significantly decreased the escalation of cocaine intake. **(D, E)** C646 significantly decreased cue-induced active nosepokes on both WD1 and WD28 (WD1: two-way RM ANOVA, genotype × nosepoke interactions: F_1, 12_ = 9.12, P = 0.01; post hoc, P < 0.01; WD28: two-way RM ANOVA, genotype × nosepoke interactions: F_1, 12_ = 17.81, P < 0.01; post hoc, P < 0.01; n = 7 per group). Data are presented as mean ± SEM. P < 0.05 was considered statistically significant. Statistical significance was determined by two-way repeated-measures (RM) ANOVA followed by Bonferroni’s multiple comparisons test.

**Figure S10.**
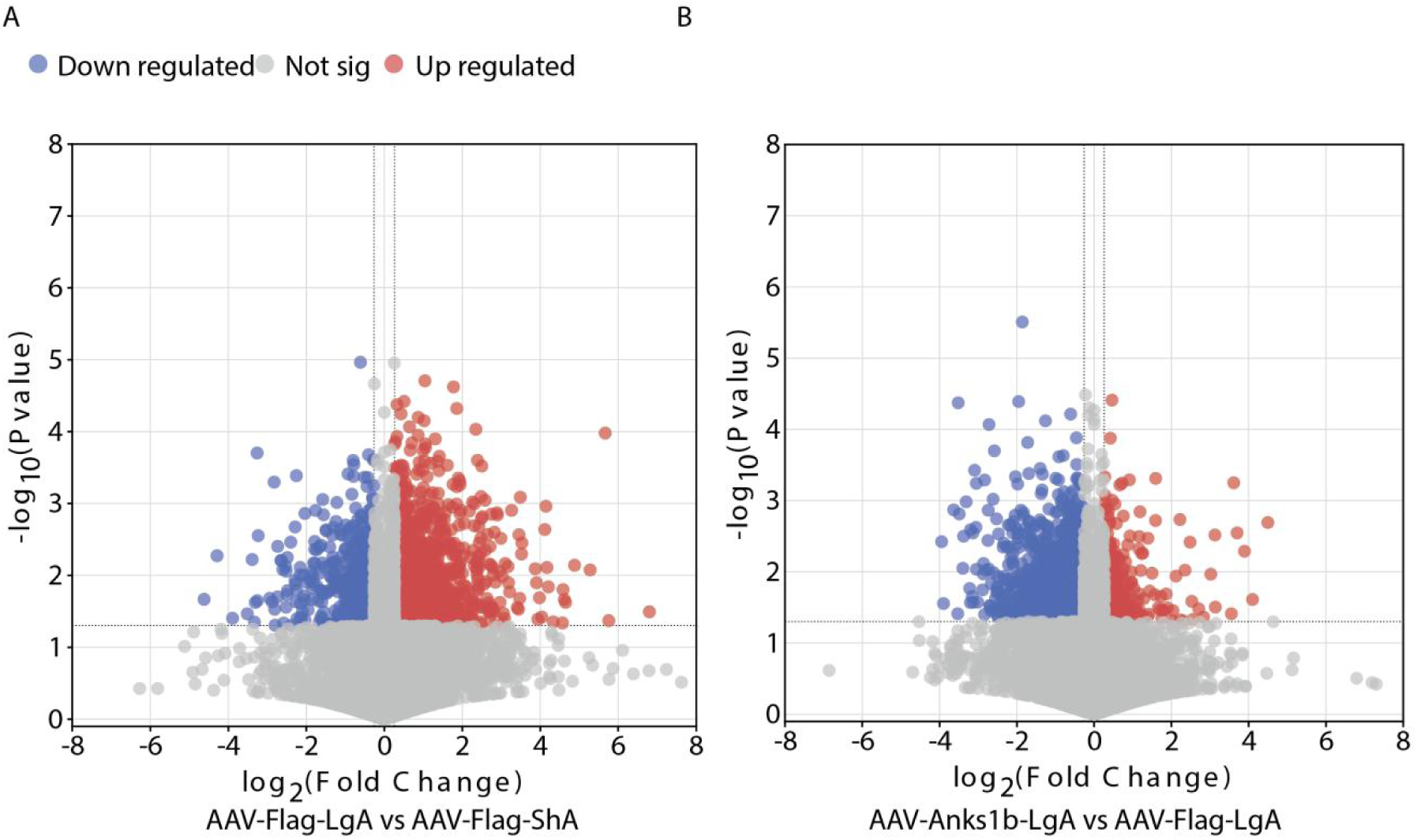
Transcriptomic analysis of gene expression changes in the Nucleus Accumbens (NAc). **(A)** Volcano plot illustrating differentially expressed genes in the NAc between the long-access (LgA) and short-access (ShA) cocaine self-administration groups. Red dots indicate significantly upregulated genes, and blue dots indicate significantly downregulated genes in the LgA group compared to the ShA group. **(B)** Volcano plot illustrating differentially expressed genes between the ANKS1B overexpression LgA group (OE-LgA) and the LgA control group. Genes downregulated by LgA are now upregulated (red dots) and genes upregulated by LgA are now downregulated (blue dots).

**Fig S11.**
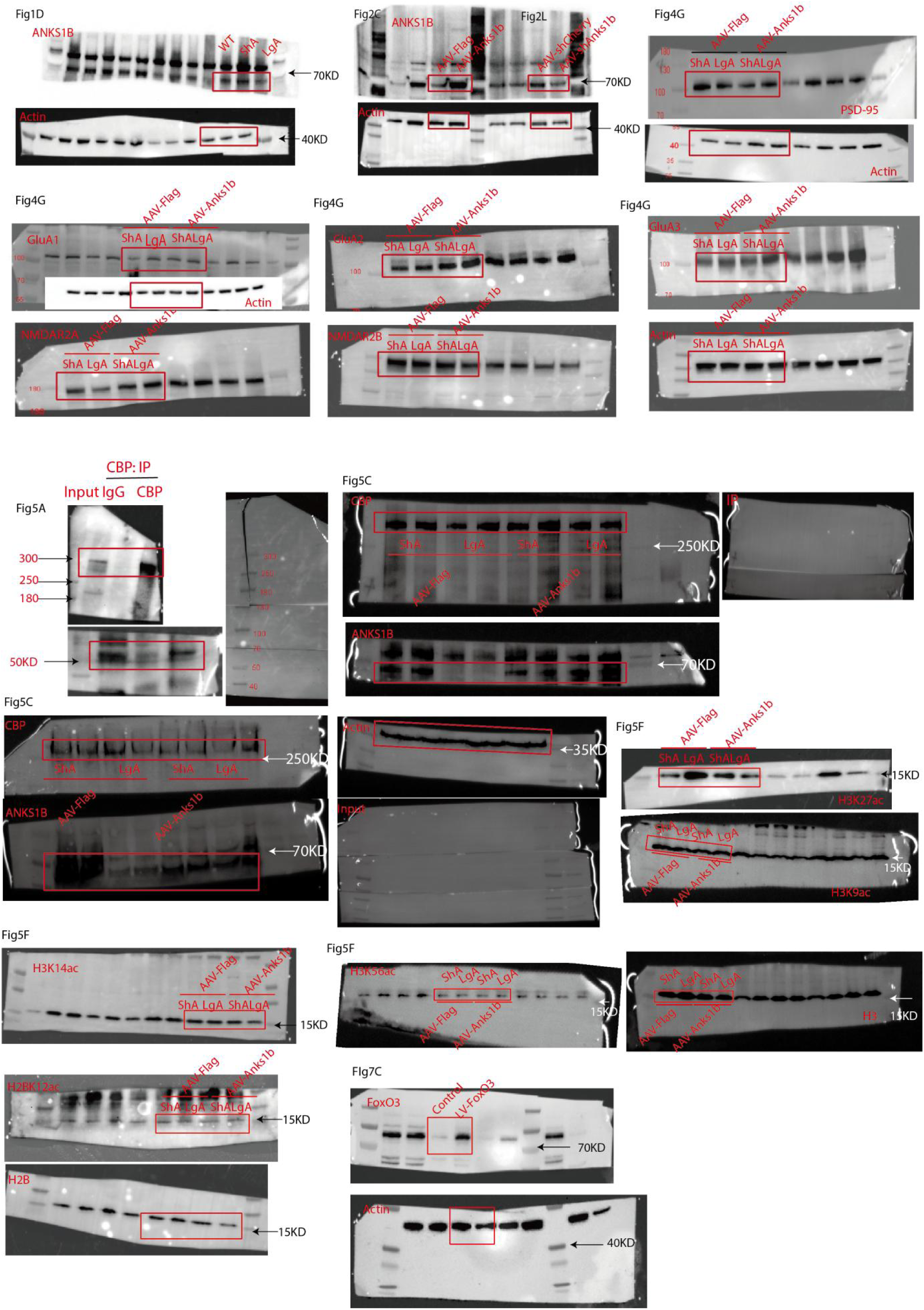
Unedited blot and gel images

---

**Table S1.**
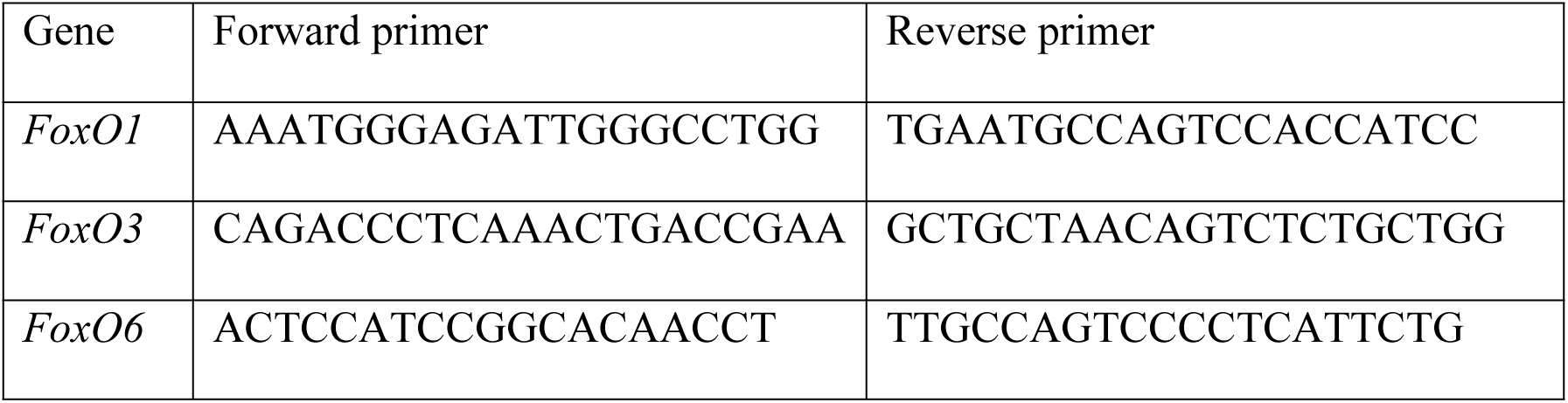
The sequences of used primers for RT-qPCR.

**Table S2.**
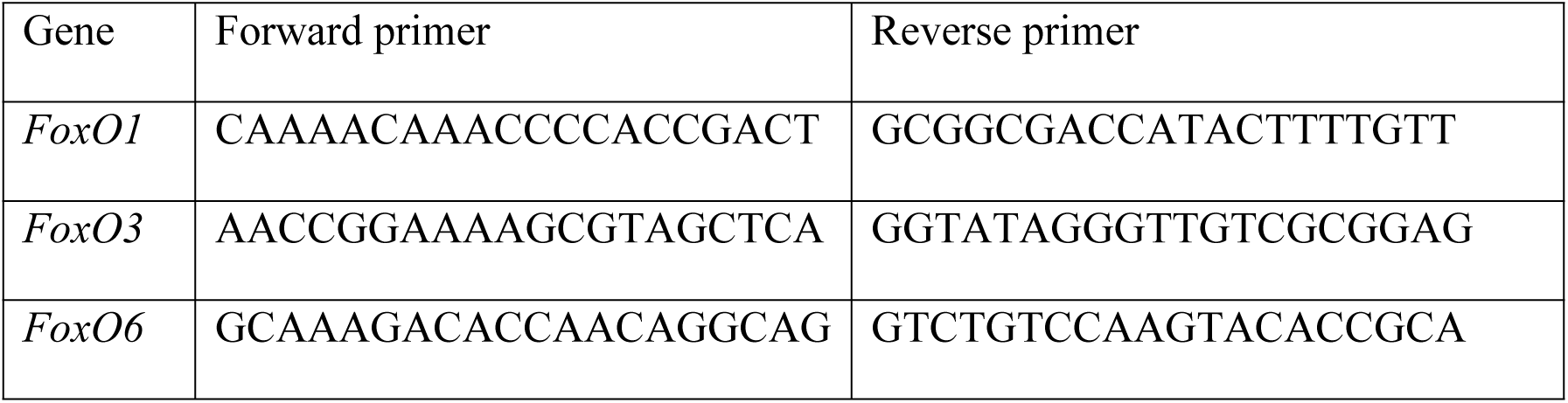
The sequences of used primers for CHIP-qPCR.

**Table S3.** Predicted confidence metrics for ANKS1B–CBP domain interaction models generated by AlphaFold3.

|  | KIX | HAT | BRO |
| --- | --- | --- | --- |
| pTM | 0.33 | 0.34 | 0.31 |
| ipTM | 0.49 | 0.70 | 0.46 |
| Ranking Score | 0.66 | 0.82 | 0.65 |

## References

1. M. Farrell, N. K. Martin, E. Stockings, et al., “Responding to global stimulant use: challenges and opportunities”, Lancet 394, no. 10209 (2019): 1652–1667. 10.1016/S0140-6736(19)32230-5

2. A. J. Butler, J. Rehm, and B. Fischer, “Health outcomes associated with crack-cocaine use: Systematic review and meta-analyses”, Drug Alcohol Depend 180, no. (2017): 401–416. 10.1016/j.drugalcdep.2017.08.036

3. A. R. Burke, J. F. DeBold, and K. A. Miczek, “CRF type 1 receptor antagonism in ventral tegmental area of adolescent rats during social defeat: prevention of escalated cocaine self-administration in adulthood and behavioral adaptations during adolescence”, Psychopharmacology (Berl) 233, no. 14 (2016): 2727–36. 10.1007/s00213-016-4336-4

4. S. H. Ahmed, and G. F. Koob, “Transition from moderate to excessive drug intake: change in hedonic set point”, Science 282, no. 5387 (1998): 298–300. 10.1126/science.282.5387.298

5. S. H. Ahmed, “The science of making drug-addicted animals”, Neuroscience 211, no. (2012): 107–25. 10.1016/j.neuroscience.2011.08.014

6. S. M. Groman, A. T. Hillmer, H. Liu, et al., “Midbrain D(3) Receptor Availability Predicts Escalation in Cocaine Self-administration”, Biol Psychiatry 88, no. 10 (2020): 767–776. 10.1016/j.biopsych.2020.02.017

7. A. U. Carbonell, C. H. Cho, J. O. Tindi, et al., “Haploinsufficiency in the ANKS1B gene encoding AIDA-1 leads to a neurodevelopmental syndrome”, Nat Commun 10, no. 1 (2019): 3529. 10.1038/s41467-019-11437-w

8. J. O. Tindi, A. E. Chávez, S. Cvejic, E. Calvo-Ochoa, P. E. Castillo, and B. A. Jordan, “ANKS1B Gene Product AIDA-1 Controls Hippocampal Synaptic Transmission by Regulating GluN2B Subunit Localization”, The Journal of Neuroscience: the Official Journal of the Society For Neuroscience 35, no. 24 (2015): 8986–96. 10.1523/jneurosci.4029-14.2015

9. A. U. Carbonell, C. Freire-Cobo, I. V. Deyneko, et al., “Comparing synaptic proteomes across five mouse models for autism reveals converging molecular similarities including deficits in oxidative phosphorylation and Rho GTPase signaling”, Front Aging Neurosci 15, no. (2023): 1152562. 10.3389/fnagi.2023.1152562

10. R. M. Younis, R. M. Taylor, P. M. Beardsley, and J. L. McClay, “The ANKS1B gene and its associated phenotypes: focus on CNS drug response”, Pharmacogenomics 20, no. 9 (2019): 669–684. 10.2217/pgs-2019-0015

11. A. Parra-Damas, and C. A. Saura, “Synapse-to-Nucleus Signaling in Neurodegenerative and Neuropsychiatric Disorders”, Biol Psychiatry 86, no. 2 (2019): 87–96. 10.1016/j.biopsych.2019.01.006

12. B. Mahjani, K. Bey, J. Boberg, and C. Burton, “Genetics of obsessive-compulsive disorder”, Psychol Med 51, no. 13 (2021): 2247–2259. 10.1017/s0033291721001744

13. E. Grünblatt, B. Oneda, A. B. Ekici, et al., “High resolution chromosomal microarray analysis in paediatric obsessive-compulsive disorder”, BMC Med Genomics 10, no. 1 (2017): 68. 10.1186/s12920-017-0299-5

14. P. J. Mulholland, B. A. Jordan, and L. J. Chandler, “Chronic ethanol up-regulates the synaptic expression of the nuclear translational regulatory protein AIDA-1 in primary hippocampal neurons”, Alcohol 46, no. 6 (2012): 569–76. 10.1016/j.alcohol.2012.04.004

15. H. Sun, J. A. Martin, C. T. Werner, et al., “BAZ1B in Nucleus Accumbens Regulates Reward-Related Behaviors in Response to Distinct Emotional Stimuli”, The Journal of Neuroscience: the Official Journal of the Society For Neuroscience 36, no. 14 (2016): 3954–61. 10.1523/jneurosci.3254-15.2016

16. S. H. Chang, Y. Sun, F. Wang, et al., “Genome-wide association meta-analyses identify novel genetic risk loci and polygenic phenotype associations for heroin, methamphetamine and alcohol dependences”, Clin Transl Med 12, no. 1 (2022): e659. 10.1002/ctm2.659

17. Y. Sun, S. Chang, Z. Liu, et al., “Identification of novel risk loci with shared effects on alcoholism, heroin, and methamphetamine dependence”, Mol Psychiatry 26, no. 4 (2021): 1152–1161. 10.1038/s41380-019-0497-y

18. E. A. Griffin, Jr., P. A. Melas, R. Zhou, et al., “Prior alcohol use enhances vulnerability to compulsive cocaine self-administration by promoting degradation of HDAC4 and HDAC5”, Sci Adv 3, no. 11 (2017): e1701682. 10.1126/sciadv.1701682

19. E. Borrelli, E. J. Nestler, C. D. Allis, and P. Sassone-Corsi, “Decoding the epigenetic language of neuronal plasticity”, Neuron 60, no. 6 (2008): 961–74. 10.1016/j.neuron.2008.10.012

20. R. R. Campbell, E. A. Kramár, L. Pham, et al., “HDAC3 Activity within the Nucleus Accumbens Regulates Cocaine-Induced Plasticity and Behavior in a Cell-Type-Specific Manner”, The Journal of Neuroscience: the Official Journal of the Society For Neuroscience 41, no. 13 (2021): 2814–2827. 10.1523/jneurosci.2829-20.2021

21. A. Rudenko, and L. H. Tsai, “Epigenetic modifications in the nervous system and their impact upon cognitive impairments”, Neuropharmacology 80, no. (2014): 70–82. 10.1016/j.neuropharm.2014.01.043

22. S. Y. Roth, J. M. Denu, and C. D. Allis, “Histone acetyltransferases”, Annu Rev Biochem 70, no. (2001): 81–120. 10.1146/annurev.biochem.70.1.81

23. M. W. Feltenstein, R. E. See, and R. A. Fuchs, “Neural Substrates and Circuits of Drug Addiction”, Cold Spring Harb Perspect Med 11, no. 4 (2021): 10.1101/cshperspect.a039628

24. B. J. Everitt, and T. W. Robbins, “Drug Addiction: Updating Actions to Habits to Compulsions Ten Years On”, Annu Rev Psychol 67, no. (2016): 23–50. 10.1146/annurev-psych-122414-033457

25. H. M. Cates, E. A. Heller, C. K. Lardner, et al., “Transcription Factor E2F3a in Nucleus Accumbens Affects Cocaine Action via Transcription and Alternative Splicing”, Biol Psychiatry 84, no. 3 (2018): 167–179. 10.1016/j.biopsych.2017.11.027

26. R. Massart, R. Barnea, Y. Dikshtein, et al., “Role of DNA methylation in the nucleus accumbens in incubation of cocaine craving”, The Journal of Neuroscience: the Official Journal of the Society For Neuroscience 35, no. 21 (2015): 8042–8058. 10.1523/JNEUROSCI.3053-14.2015

27. S. R. Cunha, and P. J. Mohler, “Ankyrin protein networks in membrane formation and stabilization”, J Cell Mol Med 13, no. 11-12 (2009): 4364–76. 10.1111/j.1582-4934.2009.00943.x

28. B. A. Jordan, B. D. Fernholz, M. Boussac, et al., “Identification and verification of novel rodent postsynaptic density proteins”, Mol Cell Proteomics 3, no. 9 (2004): 857–71. 10.1074/mcp.M400045-MCP200

29. M. Wu, Y. Zhao, J. Yang, et al., “The role of ankyrin repeat-containing proteins in epigenetic and transcriptional regulation”, Cell Death Discov 11, no. 1 (2025): 232. 10.1038/s41420-025-02519-4

30. J. Liu, B. Johnson, R. Wu, et al., “TAAR1 agonists attenuate extended-access cocaine self-administration and yohimbine-induced reinstatement of cocaine-seeking”, Br J Pharmacol 177, no. 15 (2020): 3403–3414. 10.1111/bph.15061

31. C. A. Blackwood, M. T. McCoy, B. Ladenheim, and J. L. Cadet, “Oxycodone self-administration activates the mitogen-activated protein kinase/ mitogen- and stress-activated protein kinase (MAPK-MSK) signaling pathway in the rat dorsal striatum”, Sci Rep 11, no. 1 (2021): 2567. 10.1038/s41598-021-82206-3

32. G. de Guglielmo, L. Carrette, M. Kallupi, et al., “Large-scale characterization of cocaine addiction-like behaviors reveals that escalation of intake, aversion-resistant responding, and breaking-points are highly correlated measures of the same construct”, Elife 12, no. (2024): 10.7554/eLife.90422

33. B. W. Hughes, J. L. Huebschman, E. Tsvetkov, et al., “NPAS4 supports cocaine-conditioned cues in rodents by controlling the cell type-specific activation balance in the nucleus accumbens”, Nat Commun 15, no. 1 (2024): 5971. 10.1038/s41467-024-50099-1

34. L. Caffino, F. Mottarlini, G. Targa, M. M. M. Verheij, F. Fumagalli, and J. R. Homberg, “Responsivity of serotonin transporter knockout rats to short and long access to cocaine: Modulation of the glutamate signalling in the nucleus accumbens shell”, Br J Pharmacol 179, no. 14 (2022): 3727–3739. 10.1111/bph.15823

35. S. E. Doyle, C. Ramôa, G. Garber, J. Newman, Z. Toor, and W. J. Lynch, “A shift in the role of glutamatergic signaling in the nucleus accumbens core with the development of an addicted phenotype”, Biol Psychiatry 76, no. 10 (2014): 810–5. 10.1016/j.biopsych.2014.02.005

36. C. R. Ferrario, G. Gorny, H. S. Crombag, Y. Li, B. Kolb, and T. E. Robinson, “Neural and behavioral plasticity associated with the transition from controlled to escalated cocaine use”, Biol Psychiatry 58, no. 9 (2005): 751–9. 10.1016/j.biopsych.2005.04.046

37. E. Korzus, M. G. Rosenfeld, and M. Mayford, “CBP histone acetyltransferase activity is a critical component of memory consolidation”, Neuron 42, no. 6 (2004): 961–72. 10.1016/j.neuron.2004.06.002

38. Y. Funahashi, A. Ariza, R. Emi, et al., “Phosphorylation of Npas4 by MAPK Regulates Reward-Related Gene Expression and Behaviors”, Cell Rep 29, no. 10 (2019): 3235–3252.e9. 10.1016/j.celrep.2019.10.116

39. S. Kosol, S. Contreras-Martos, A. Piai, et al., “Interaction between the scaffold proteins CBP by IQGAP1 provides an interface between gene expression and cytoskeletal activity”, Sci Rep 10, no. 1 (2020): 5753. 10.1038/s41598-020-62069-w

40. V. V. Ogryzko, R. L. Schiltz, V. Russanova, B. H. Howard, and Y. Nakatani, “The transcriptional coactivators p300 and CBP are histone acetyltransferases”, Cell 87, no. 5 (1996): 953–9. 10.1016/s0092-8674(00)82001-2

41. J. S. Guan, S. J. Haggarty, E. Giacometti, et al., “HDAC2 negatively regulates memory formation and synaptic plasticity”, Nature 459, no. 7243 (2009): 55–60. 10.1038/nature07925

42. E. Yazici, and J. McIntyre, “The complex network of p300/CBP regulation: Interactions, posttranslational modifications, and therapeutic implications”, J Biol Chem 301, no. 11 (2025): 110715. 10.1016/j.jbc.2025.110715

43. F. Tie, R. Banerjee, C. Fu, C. A. Stratton, M. Fang, and P. J. Harte, “Polycomb inhibits histone acetylation by CBP by binding directly to its catalytic domain”, Proc Natl Acad Sci U S A 113, no. 6 (2016): E744–53. 10.1073/pnas.1515465113

44. D. Cheng, Z. Dong, P. Lin, G. Shen, and Q. Xia, “Transcriptional Activation of Ecdysone-Responsive Genes Requires H3K27 Acetylation at Enhancers”, Int J Mol Sci 23, no. 18 (2022): 10.3390/ijms231810791

45. J. Chen, L. Wang, G. G. Tian, X. Wang, X. Li, and J. Wu, “Metformin Promotes Proliferation of Mouse Female Germline Stem Cells by Histone Acetylation Modification of Traf2”, Stem Cell Rev Rep 19, no. 7 (2023): 2329–2340. 10.1007/s12015-023-10575-5

46. D. L. Gerrard, J. R. Boyd, G. S. Stein, V. X. Jin, and S. Frietze, “Disruption of Broad Epigenetic Domains in PDAC Cells by HAT Inhibitors”, Epigenomes 3, no. 2 (2019): 10.3390/epigenomes3020011

47. S. Y. Kim, and A. E. Webb, “Neuronal functions of FOXO/DAF-16”, Nutr Healthy Aging 4, no. 2 (2017): 113–126. 10.3233/nha-160009

48. E. E. Santo, and J. Paik, “FOXO in Neural Cells and Diseases of the Nervous System”, Curr Top Dev Biol 127, no. (2018): 105–118. 10.1016/bs.ctdb.2017.10.002

49. T. Rana, T. Behl, A. Sehgal, et al., “Elucidating the Possible Role of FoxO in Depression”, Neurochem Res 46, no. 11 (2021): 2761–2775. 10.1007/s11064-021-03364-4

50. E. H. Jang, and S. A. Kim, “Long-Term Epigenetic Regulation of Foxo3 Expression in Neonatal Valproate-Exposed Rat Hippocampus with Sex-Related Differences”, Int J Mol Sci 25, no. 10 (2024): 10.3390/ijms25105287

51. S. Salcher, G. Spoden, J. Hagenbuchner, et al., “A drug library screen identifies Carbenoxolone as novel FOXO inhibitor that overcomes FOXO3-mediated chemoprotection in high-stage neuroblastoma”, Oncogene 39, no. 5 (2020): 1080–1097. 10.1038/s41388-019-1044-7

52. S. K. Son, J. S. Moon, D. W. Yang, et al., “Role of FOXO3a in LPS-induced inflammatory conditions in human dental pulp cells”, J Oral Biosci 67, no. 1 (2025): 100614. 10.1016/j.job.2025.100614

53. B. E. Schmeichel, A. Matzeu, P. Koebel, et al., “Knockdown of hypocretin attenuates extended access of cocaine self-administration in rats”, Neuropsychopharmacology 43, no. 12 (2018): 2373–2382. 10.1038/s41386-018-0054-4

54. C. Nicolas, C. Tauber, F. X. Lepelletier, et al., “Longitudinal Changes in Brain Metabolic Activity after Withdrawal from Escalation of Cocaine Self-Administration”, Neuropsychopharmacology 42, no. 10 (2017): 1981–1990. 10.1038/npp.2017.109

55. C. A. Winstanley, Q. LaPlant, D. E. Theobald, et al., “DeltaFosB induction in orbitofrontal cortex mediates tolerance to cocaine-induced cognitive dysfunction”, The Journal of Neuroscience: the Official Journal of the Society For Neuroscience 27, no. 39 (2007): 10497–507. 10.1523/jneurosci.2566-07.2007

56. K. Vaillancourt, C. Ernst, D. Mash, and G. Turecki, “DNA Methylation Dynamics and Cocaine in the Brain: Progress and Prospects”, Genes (Basel) 8, no. 5 (2017): 10.3390/genes8050138

57. K. Vaillancourt, G. G. Chen, L. Fiori, et al., “Methylation of the tyrosine hydroxylase gene is dysregulated by cocaine dependence in the human striatum”, iScience 24, no. 10 (2021): 103169. 10.1016/j.isci.2021.103169

58. K. Anier, M. Urb, K. Kipper, et al., “Cocaine-induced epigenetic DNA modification in mouse addiction-specific and non-specific tissues”, Neuropharmacology 139, no. (2018): 13–25. 10.1016/j.neuropharm.2018.06.036

59. M. Urb, K. Niinep, T. Matsalu, et al., “The role of DNA methyltransferase activity in cocaine treatment and withdrawal in the nucleus accumbens of mice”, Addict Biol 25, no. 1 (2020): e12720. 10.1111/adb.12720

60. R. Nudel, R. Zetterberg, N. Hemager, et al., “A family-based study of genetic and epigenetic effects across multiple neurocognitive, motor, social-cognitive and social-behavioral functions”, Behav Brain Funct 18, no. 1 (2022): 14. 10.1186/s12993-022-00198-0

61. L. A. Papale, A. Madrid, S. Li, and R. S. Alisch, “Early-life stress links 5-hydroxymethylcytosine to anxiety-related behaviors”, Epigenetics 12, no. 4 (2017): 264–276. 10.1080/15592294.2017.1285986

62. L. A. Papale, S. Li, A. Madrid, et al., “Sex-specific hippocampal 5-hydroxymethylcytosine is disrupted in response to acute stress”, Neurobiology of Disease 96, no. (2016): 54–66. 10.1016/j.nbd.2016.08.014

63. C. T. Watson, H. Szutorisz, P. Garg, et al., “Genome-Wide DNA Methylation Profiling Reveals Epigenetic Changes in the Rat Nucleus Accumbens Associated With Cross-Generational Effects of Adolescent THC Exposure”, Neuropsychopharmacology 40, no. 13 (2015): 2993–3005. 10.1038/npp.2015.155

64. J. Bourne, and K. M. Harris, “Do thin spines learn to be mushroom spines that remember?”, Curr Opin Neurobiol 17, no. 3 (2007): 381–6. 10.1016/j.conb.2007.04.009

65. F. Kasanetz, V. Deroche-Gamonet, N. Berson, et al., “Transition to addiction is associated with a persistent impairment in synaptic plasticity”, Science 328, no. 5986 (2010): 1709–12. 10.1126/science.1187801

66. I. Smaga, M. Sanak, and M. Filip, “Cocaine-induced Changes in the Expression of NMDA Receptor Subunits”, Curr Neuropharmacol 17, no. 11 (2019): 1039–1055. 10.2174/1570159x17666190617101726

67. T. Su, Y. Lu, C. Fu, Y. Geng, and Y. Chen, “GluN2A mediates ketamine-induced rapid antidepressant-like responses”, Nat Neurosci 26, no. 10 (2023): 1751–1761. 10.1038/s41593-023-01436-y

68. Q. Q. Li, J. Chen, P. Hu, et al., “Enhancing GluN2A-type NMDA receptors impairs long-term synaptic plasticity and learning and memory”, Mol Psychiatry 27, no. 8 (2022): 3468–3478. 10.1038/s41380-022-01579-7

69. J. Feng, and E. J. Nestler, “Epigenetic mechanisms of drug addiction”, Curr Opin Neurobiol 23, no. 4 (2013): 521–8. 10.1016/j.conb.2013.01.001

70. M. J. Bannon, M. M. Johnson, S. K. Michelhaugh, et al., “A molecular profile of cocaine abuse includes the differential expression of genes that regulate transcription, chromatin, and dopamine cell phenotype”, Neuropsychopharmacology 39, no. 9 (2014): 2191–9. 10.1038/npp.2014.70

71. C. T. Werner, S. Mitra, J. A. Martin, et al., “Ubiquitin-proteasomal regulation of chromatin remodeler INO80 in the nucleus accumbens mediates persistent cocaine craving”, Sci Adv 5, no. 10 (2019): eaay0351. 10.1126/sciadv.aay0351

72. S. H. Ahmed, P. J. Kenny, G. F. Koob, and A. Markou, “Neurobiological evidence for hedonic allostasis associated with escalating cocaine use”, Nat Neurosci 5, no. 7 (2002): 625–6. 10.1038/nn872

73. A. S. Koijam, K. D. Singh, B. S. Nameirakpam, R. Haobam, and Y. Rajashekar, “Drug addiction and treatment: An epigenetic perspective”, Biomed Pharmacother 170, no. (2024): 115951. 10.1016/j.biopha.2023.115951

74. G. C. Sartor, S. K. Powell, S. P. Brothers, and C. Wahlestedt, “Epigenetic Readers of Lysine Acetylation Regulate Cocaine-Induced Plasticity”, The Journal of Neuroscience: the Official Journal of the Society For Neuroscience 35, no. 45 (2015): 15062–72. 10.1523/jneurosci.0826-15.2015

75. J. Cheng, Z. He, Q. Chen, et al., “Histone modifications in cocaine, methamphetamine and opioids”, Heliyon 9, no. 6 (2023): e16407. 10.1016/j.heliyon.2023.e16407

76. A. A. Levine, Z. Guan, A. Barco, S. Xu, E. R. Kandel, and J. H. Schwartz, “CREB-binding protein controls response to cocaine by acetylating histones at the fosB promoter in the mouse striatum”, Proc Natl Acad Sci U S A 102, no. 52 (2005): 19186–91. 10.1073/pnas.0509735102

77. M. Malvaez, E. Mhillaj, D. P. Matheos, M. Palmery, and M. A. Wood, “CBP in the nucleus accumbens regulates cocaine-induced histone acetylation and is critical for cocaine-associated behaviors”, The Journal of Neuroscience: the Official Journal of the Society For Neuroscience 31, no. 47 (2011): 16941–8. 10.1523/jneurosci.2747-11.2011

78. D. Pasini, M. Malatesta, H. R. Jung, et al., “Characterization of an antagonistic switch between histone H3 lysine 27 methylation and acetylation in the transcriptional regulation of Polycomb group target genes”, Nucleic Acids Res 38, no. 15 (2010): 4958–69. 10.1093/nar/gkq244

79. A. Zhang, P. L. Yeung, C. W. Li, et al., “Identification of a novel family of ankyrin repeats containing cofactors for p160 nuclear receptor coactivators”, J Biol Chem 279, no. 32 (2004): 33799–805. 10.1074/jbc.M403997200

80. D. Gallagher, A. Voronova, M. A. Zander, et al., “Ankrd11 is a chromatin regulator involved in autism that is essential for neural development”, Dev Cell 32, no. 1 (2015): 31–42. 10.1016/j.devcel.2014.11.031

81. T. A. McKinsey, K. Kuwahara, S. Bezprozvannaya, and E. N. Olson, “Class II histone deacetylases confer signal responsiveness to the ankyrin-repeat proteins ANKRA2 and RFXANK”, Mol Biol Cell 17, no. 1 (2006): 438–47. 10.1091/mbc.e05-07-0612

82. S. M. Truhlar, J. W. Torpey, and E. A. Komives, “Regions of IkappaBalpha that are critical for its inhibition of NF-kappaB.DNA interaction fold upon binding to NF-kappaB”, Proc Natl Acad Sci U S A 103, no. 50 (2006): 18951–6. 10.1073/pnas.0605794103

83. S. Majumder, G. Zhu, X. Xu, et al., “G-Protein-Coupled Receptor-2-Interacting Protein-1 Controls Stalk Cell Fate by Inhibiting Delta-like 4-Notch1 Signaling”, Cell Rep 17, no. 10 (2016): 2532–2541. 10.1016/j.celrep.2016.11.017

84. K. Maiese, Z. Z. Chong, and Y. C. Shang, ””Sly as a FOXO”: new paths with Forkhead signaling in the brain”, Curr Neurovasc Res 4, no. 4 (2007): 295–302. 10.2174/156720207782446306

85. T. He, C. Han, C. Liu, et al., “Dopamine D1 receptors mediate methamphetamine-induced dopaminergic damage: involvement of autophagy regulation via the AMPK/FOXO3A pathway”, Psychopharmacology (Berl) 239, no. 3 (2022): 951–964. 10.1007/s00213-022-06097-6

86. V. Kumaresan, Y. Lim, P. Juneja, et al., “Abstinence from Escalation of Cocaine Intake Changes the microRNA Landscape in the Cortico-Accumbal Pathway”, Biomedicines 11, no. 5 (2023): 10.3390/biomedicines11051368

87. Y. Wang, L. Yang, H. Zhou, K. Zhang, and M. Zhao, “Identification of miRNA-mediated gene regulatory networks in L-methionine exposure counteracts cocaine-conditioned place preference in mice”, Front Genet 13, no. (2022): 1076156. 10.3389/fgene.2022.1076156

88. H. F. Lu, W. Xiao, S. L. Deng, et al., “Activation of AMPK-dependent autophagy in the nucleus accumbens opposes cocaine-induced behaviors of mice”, Addict Biol 25, no. 2 (2020): e12736. 10.1111/adb.12736

89. D. Ferguson, N. Shao, E. Heller, et al., “SIRT1-FOXO3a regulate cocaine actions in the nucleus accumbens”, The Journal of Neuroscience: the Official Journal of the Society For Neuroscience 35, no. 7 (2015): 3100–11. 10.1523/jneurosci.4012-14.2015

90. C. N. McLaughlin, and H. T. Broihier, “Keeping Neurons Young and Foxy: FoxOs Promote Neuronal Plasticity”, Trends Genet 34, no. 1 (2018): 65–78. 10.1016/j.tig.2017.10.002

91. A. E. Webb, E. A. Pollina, T. Vierbuchen, et al., “FOXO3 shares common targets with ASCL1 genome-wide and inhibits ASCL1-dependent neurogenesis”, Cell Rep 4, no. 3 (2013): 477–91. 10.1016/j.celrep.2013.06.035

92. Y. X. Xue, Y. X. Luo, P. Wu, et al., “A memory retrieval-extinction procedure to prevent drug craving and relapse”, Science 336, no. 6078 (2012): 241–5. 10.1126/science.1215070

93. L. Kokarovtseva, T. Jaciw-Zurakiwsky, R. Mendizabal Arbocco, M. V. Frantseva, and J. L. Perez Velazquez, “Excitability and gap junction-mediated mechanisms in nucleus accumbens regulate self-stimulation reward in rats”, Neuroscience 159, no. 4 (2009): 1257–63. 10.1016/j.neuroscience.2009.01.065

94. S. A. Maddox, C. S. Watts, and G. E. Schafe, “p300/CBP histone acetyltransferase activity is required for newly acquired and reactivated fear memories in the lateral amygdala”, Learn Mem 20, no. 2 (2013): 109–19. 10.1101/lm.029157.112

95. W. Chen, Y. Zhang, J. Liang, et al., “Disrupting astrocyte-neuron lactate transport prevents cocaine seeking after prolonged withdrawal”, Sci Adv 9, no. 43 (2023): eadi4462. 10.1126/sciadv.adi4462

96. Z. Yu, W. Chen, L. Zhang, et al., “Gut-derived bacterial LPS attenuates incubation of methamphetamine craving via modulating microglia”, Brain Behav Immun 111, no. (2023): 101–115. 10.1016/j.bbi.2023.03.027

97. Z. Yu, Y. Han, D. Hu, et al., “Neurocan regulates vulnerability to stress and the anti-depressant effect of ketamine in adolescent rats”, Mol Psychiatry 27, no. 5 (2022): 2522–2532. 10.1038/s41380-022-01495-w

98. Z. Yu, N. Chen, D. Hu, et al., “Decreased Density of Perineuronal Net in Prelimbic Cortex Is Linked to Depressive-Like Behavior in Young-Aged Rats”, Front Mol Neurosci 13, no. (2020): 4. 10.3389/fnmol.2020.00004

99. R. C. Gruber, G. S. Wirak, A. S. Blazier, et al., “BTK regulates microglial function and neuroinflammation in human stem cell models and mouse models of multiple sclerosis”, Nat Commun 15, no. 1 (2024): 10116. 10.1038/s41467-024-54430-8

100. E. M. Glaser, and H. Van der Loos, “Analysis of thick brain sections by obverse-reverse computer microscopy: application of a new, high clarity Golgi-Nissl stain”, J Neurosci Methods 4, no. 2 (1981): 117–25. 10.1016/0165-0270(81)90045-5

101. Y. Han, Y. Luo, J. Sun, et al., “AMPK Signaling in the Dorsal Hippocampus Negatively Regulates Contextual Fear Memory Formation”, Neuropsychopharmacology 41, no. 7 (2016): 1849–64. 10.1038/npp.2015.355

